# GLI transcriptional repression is inert prior to Hedgehog pathway activation

**DOI:** 10.1101/2021.06.29.450392

**Authors:** Rachel K. Lex, Weiqiang Zhou, Zhicheng Ji, Kristin N. Falkenstein, Kaleigh E. Schuler, Kathryn E. Windsor, Joseph D. Kim, Hongkai Ji, Steven A. Vokes

**Affiliations:** Department of Molecular Biosciences, University of Texas at Austin, 100 E 24th Street Stop A5000, Austin, TX 78712 USA; Department of Biostatistics, Johns Hopkins Bloomberg School of Public Health, 615 North Wolfe Street, Room E3638, Baltimore, MD 21205, USA; Department of Biostatistics and Bioinformatics, Duke University School of Medicine, 2424 Erwin Road, Suite 1102 Hock Plaza Box 2721 Durham, NC 27710

**Author notes:** Corresponding author Phone: -232-8359.

**Keywords:** Hedgehog signaling, GLI, GLI3, transcriptional repression, chromatin, limb bud, limb development, pre-patterning, cilia

## Abstract

In the absence of Hedgehog (HH) signaling, GLI proteins are post-translationally modified within cilia into transcriptional repressors that subsequently prevent sub-threshold activation of HH target genes. GLI repression is presumably important for preventing precocious expression of target genes before the onset of HH pathway activation, a presumption that underlies the pre-patterning model of anterior-posterior limb polarity. Here, we report that GLI3 repressor is abundant and binds to target genes in early limb development. However, contrary to expectations, GLI3 repression neither regulates the activity of GLI enhancers nor expression of HH target genes as it does after HH signaling has been established. Within the cilia, the transition to active GLI repression is accompanied by increases in axonemal GLI3 localization, possibly signifying altered GLI3 processing. Together, our results demonstrate that GLI3 repression does not prevent precocious activation of HH target genes, or have a pre-patterning role in regulating anterior-posterior limb polarity.

## Introduction

The Hedgehog (HH) signaling pathway is one of the major developmental regulators of tissue-specific development and differentiation. GLI proteins mediate transcriptional responses to the pathway in a strikingly cell-specific fashion. In the absence of HH pathway activation, GLI transcriptional repressors (GLI-R) prevent the activation of HH target genes, while upon exposure to HH, GLI proteins undergo alternative post-translational processing into transcriptional activators (GLI-A) (Aza-Blanc et al., 1997; Méthot and Basler, 1999; Panman et al., 2006; Wang et al., 2000; Niewiadomski et al., 2014). The processing of GLI-A and GLI-R are both directly dependent on processes localized within the primary cilium and defects in ciliary components lead to characteristic misregulation of HH target genes in a large group of birth defects termed ciliopathies (reviewed in Ho and Stearns, 2021). While GLI-A directly activates a subset of HH targets, most genes are ‘de-repressed’ by the loss of GLI-R alone; they do not require GLI-A-mediated activation for their expression. The importance of de-repression is exemplified in the developing limb where the phenotype of *Sonic Hedgehog (Shh)* null limb buds is dramatically improved in *Shh;Gli3* mutants (Lewandowski et al., 2015; Litingtung et al., 2002; te Welscher et al., 2002). In this context, the loss of GLI3-mediated repression, even in the absence of pathway activation, is sufficient to restore expression of most GLI target genes and many aspects of limb growth and patterning.

GLI3 represses transcription at least in part by epigenetically regulating a subset of its own enhancers. Its properties include reduced enrichment levels of the active enhancer mark H3K27ac and reduced chromatin accessibility at a subset of HH-responsive GLI3 binding regions (GBRs), termed HH-responsive GBRs, that likely mediate the majority of HH-specific transcription (Lex et al., 2020). As a regulator of tissue patterning, GLI-R spatially and temporally restricts expression of HH targets, preventing sub-threshold activation of the pathway in HH-responsive tissues (Aza-Blanc et al., 1997; Balaskas et al., 2012; Parker et al., 2011). Consequently, GLI transcriptional repression has primarily been studied in tissues with ongoing HH signaling. Although never experimentally addressed, it is widely assumed that GLI-R also regulates the expression of its target genes prior to the initiation of HH signaling. Before pathway activation, GLI3 has been proposed to have a pre-patterning role in the very early limb by repressing *Hand2*, thus limiting its expression to the posterior limb bud where it is required to activate *Shh* expression and establishing anterior-posterior polarity of the limb (Galli et al., 2010; Litingtung et al., 2002; Osterwalder et al., 2014; Vokes et al., 2008; te Welscher et al., 2002; Zhulyn et al., 2014; reviewed in Zuniga and Zeller, 2020). Curiously, *Hand2* and *Gli3* are co-expressed in early limb buds, prior to their segregation into distinct posterior and anterior domains (Osterwalder et al., 2014), raising the question of whether GLI3-R is capable of repressing *Hand2* at this time.

We initially hypothesized that GLI3-mediated repression would be important prior to HH expression for preventing premature activation of target genes by reducing H3K27ac enrichment at enhancers. Despite GLI3 binding to a majority of the same sites it occupies in the post-HH limb bud, it does not regulate activation of its enhancers or their chromatin accessibility until after the initiation of HH signaling. In addition, GLI target genes are not upregulated in limb buds lacking GLI3 as they are after the initiation of HH signaling, even though a subset of these genes appear competent to be repressed at this stage. Counter to our initial hypothesis, we conclude that the GLI3-R isoform is transcriptionally inert in early limb buds, a finding that is incompatible with the pre-patterning model for limb polarity. Interestingly, most genes subject to later GLI3 mediated repression are expressed in pre-HH limb buds, suggesting that rather than preventing their activation, GLI3 repression later regulates their spatial expression. Overall, this work demonstrates that GLI3-mediated repression of target genes is not a default state, but rather, GLI repression is established during limb development at a time point after HH induction.

## Results

### GLI3 repressor is abundant and binds to most regions prior to HH induction

To understand if GLI repression is established prior to HH signaling we first defined when the pathway is activated during limb development. The earliest detection of the canonical HH target gene *Gli1* was at 24 somites (24S), where 58% of embryos at this stage had detectable *Gli1* expression, while *Shh* was not detected until 25S (86%) (Figure 1A,B; Figure S1A,B). As 24S was the earliest detection of *Gli1* expression, we defined that stage as the onset of HH signaling and the “pre-HH” window as 21-23S, corresponding to embryonic day 9.25 (E9.25), a stage slightly earlier than previous reports (Figure 1A) (Charite et al., 2000; Lewis et al., 2001; Zhu et al., 2008).

**Figure 1.**
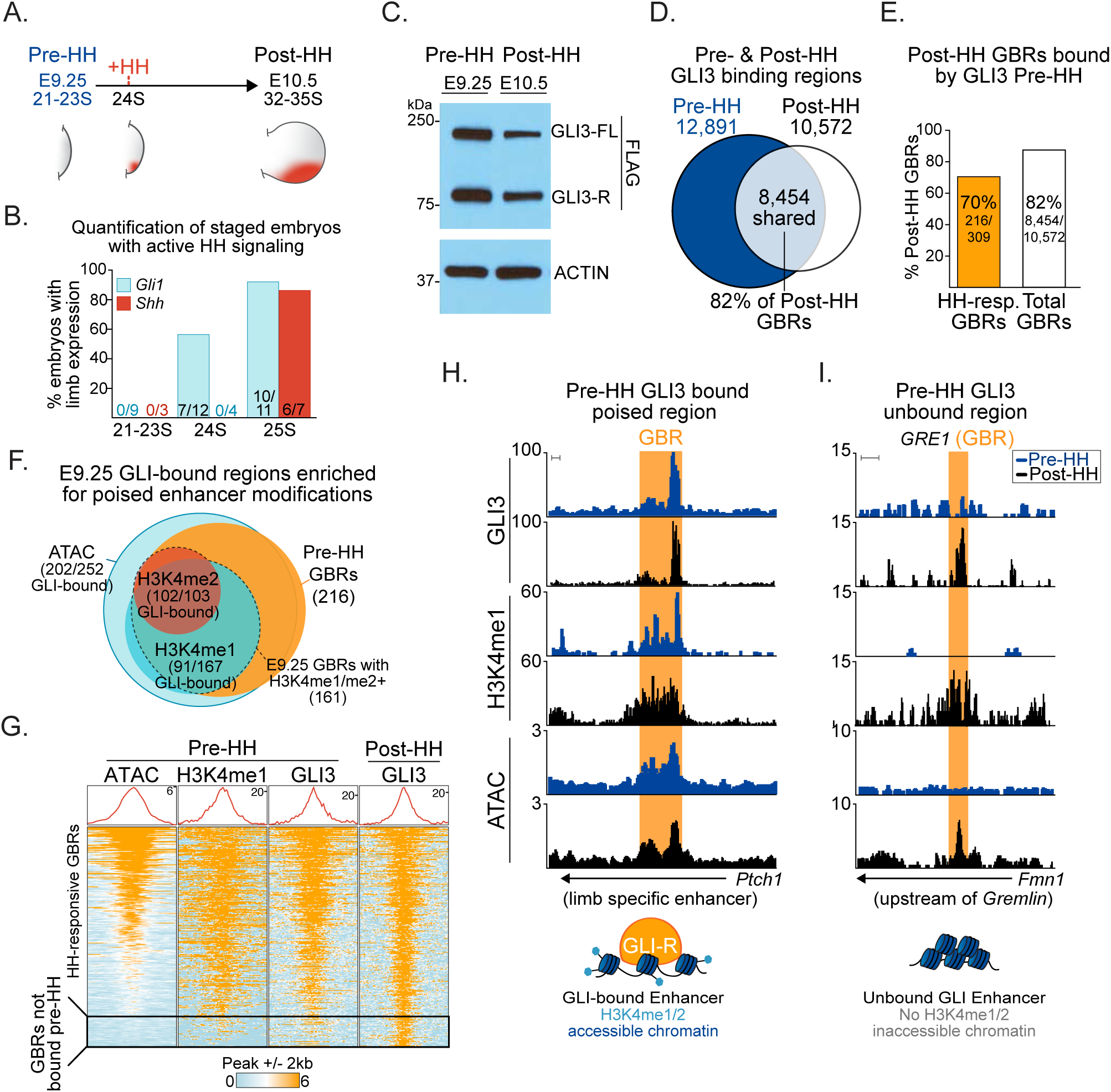
GLI3 binds to poised, accessible chromatin prior to HH signaling. A. schematic for HH induction timeline during limb development. B. Quantification of 21-25S embryos with expression of *Gli1* and *Shh* assayed by in situ hybridization. C. Representative Western blot (n=3) of endogenous GLI3^FLAG^ protein in the limb bud pre-(21-23S) and post-HH signaling (32-35S). D. Venn diagram of all pre- and post-HH identified GLI3 CUT&RUN called peaks. E. Percentage of E10.5 GLI3-bound regions also bound at E9.25, before HH signaling, for all E10.5 identified GBRs and E10.5 HH-responsive GBRs. F. Venn diagram of HH-responsive GBRs enriched for poised enhancer marks and bound by GLI3 at E9.25. Poised enhancer modifications identified by H3K4me1 CUT&Tag (n=3), H3K4me2 ChIP-seq (n=2), ATAC-seq (n=2). G. Heatmap of GLI3 enrichment pre- and post-HH signaling and enrichment of poised enhancer marks pre-HH signaling. Many regions lacking GLI3 also lack enrichment of poised marks. H. Example of a HH-responsive GBR (orange shading) that is poised, accessible, and bound by GLI3 prior to HH signaling at a limb-specific *Ptch1* enhancer (Lopez-Rios et al., 2014). I. Example of a validated HH-responsive GBR *GRE1*, that regulates the GLI3 target *Gremlin* (Li et al., 2014), that is inaccessible, lacks H3K4me1 and GLI3 binding at E9.25. Scale bars =1kb. See also Figure S1, Supplemental File 1,2.

Confirming previous findings, GLI3-R was expressed at comparable levels at both pre- and post-HH stages (Figure 1C) (Osterwalder et al., 2014). We then compared endogenous GLI3^FLAG^ binding using CUT&RUN (Skene and Henikoff, 2017) in pre-HH (E9.25, 21-23S) and E10.5 (32-35S) limb buds, a post-HH time frame when HH signaling is firmly established but before there are morphological changes in *Shh^-/-^* limb buds (Probst et al., 2011). In pre-HH limb buds, GLI3 bound to most regions that were bound in post-HH limb buds (82%; Figure 1D). Previously, we identified a group of HH-responsive GLI3 binding regions that have reduced H3K27ac in the absence of HH signaling (constitutive GLI repression) (Lex et al., 2020)(Figure S1C-E). Since this subset of GBRs seems to regulate most HH-responsive gene expression, we reasoned that these regions might be especially important to repress before HH signaling to prevent premature activation of enhancers. While the majority of HH-responsive GBRs (70%) were bound by GLI3 prior to HH induction, nearly a third were not bound, suggesting that GLI3 does not initially regulate a subset of HH-responsive GBRs (Figure 1E).

### GLI3 preferentially binds to poised, accessible enhancers

We sought to determine why GLI3 bound to only a portion of regions in the pre-HH limb and hypothesized that GLI3 may preferentially bind to poised enhancers. We defined poised enhancers as ATAC-seq accessible and enriched for either H3K4me1 or H3K4me2, where H3K4me1 is enriched at promoter-proximal and distal regions, while H3K4me2 is more commonly found promoter proximally (Ernst et al., 2011; Pekowska et al., 2011; Wang et al., 2014). Most HH-responsive GBRs that were accessible and enriched for H3K4me2 by E10.5 were already accessible (89%) and enriched for H3K4me2 (98%) in the early limb (Figure S1F,G). In contrast, of the E10.5 HH-responsive GBRs with H3K4me1, only 65% of them were enriched for H3K4me1 by E9.25 (Figure S1G). We then asked if GLI3 preferentially bound to accessible, poised regions. Consistent with this scenario, most regions bound by GLI3 in pre-HH limbs were accessible (93%) and enriched for poised enhancer modifications (75%) (Figure 1F,G; Figure S1G). These included a defined limb-specific distal enhancer for *Ptch1* (Lopez-Rios et al., 2014) (Figure 1H). While nearly half of the regions not yet bound by GLI3 in the early limb overlapped with called ATAC-seq peaks (Figure S1H), they were less accessible than GLI bound enhancers (Figure 1G). Unbound HH-responsive GBRs also generally lacked enrichment of poised enhancer marks, H3K4me1 and H3K4me2 (Figure 1G,I; Figure S1H). As many distal regions lacked H3K4me1 enrichment prior to HH induction, we observed a slight preference for GLI3 to be bound to promoter proximal regions (Figure S1I). This is exemplified by the distal limb enhancer, *GRE1* that helps regulate the HH target *Gremlin*, which is among the inaccessible regions that lack H3K4me1 enrichment and are not bound by GLI3 at E9.25 (Figure 1I). This finding is consistent with previous reports demonstrating that *GRE1* does not have enhancer activity until E10 (31-32S) (Li et al., 2014). We conclude that the majority of GLI enhancers have accessible chromatin in pre-HH limb buds. In addition, we indicate that GLI3 preferentially binds to poised enhancers, providing tissue-specific control of repression.

### HH-responsive GBRs have low levels of H3K27ac and are enriched with HDACs, before activation of the HH pathway

If GLI3 represses enhancers in the early limb as it does at E10.5, then HH-responsive GBRs, which have reduced H3K27ac in E10.5 *Shh^-/-^* limb buds, should have reduced acetylation at E9.25 before HH signaling initiates. In agreement with this scenario, there was a significant reduction in H3K27ac enrichment at HH-responsive GBRs in pre-HH compared to post-HH limb buds (Figure 2A-C). In addition, only 39% (121/309) of HH-responsive GBRs have called H3K27ac peaks in the pre-HH limb, compared to 86% (6385/7382) of all GBRs that are acetylated in the early limb bud. The overall reduction in H3K27ac enrichment at HH-responsive GBRs in the early limb bud supports the possibility that GLI3 could be actively repressing enhancers at this time to prevent premature activation.

**Figure 2.**
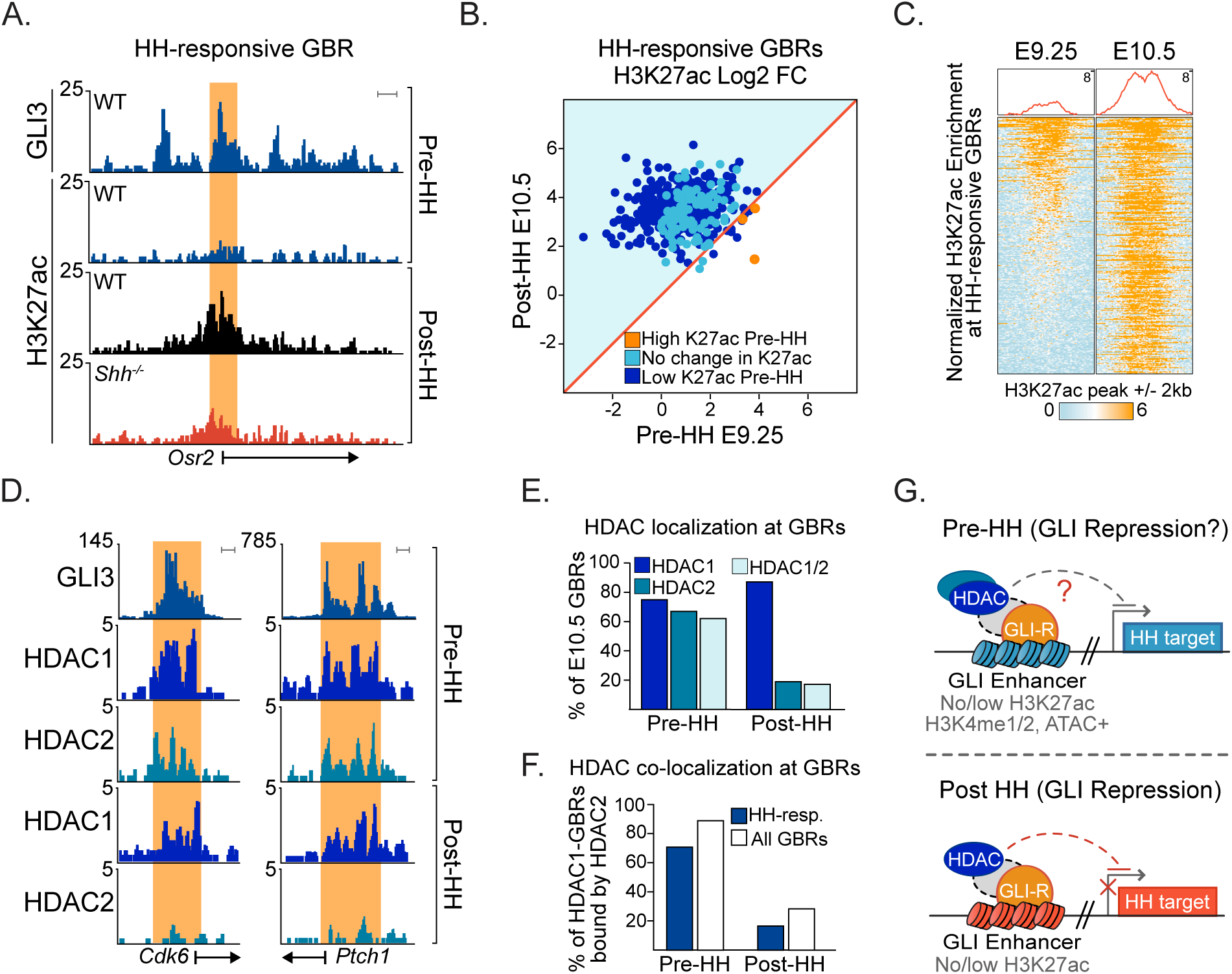
GLI3 binding regions have reduced enrichment of H3K27ac and are enriched with HDACs in pre-HH limb buds. A. H3K27ac ChIP-seq tracks showing a E9.25 (21-23S) GLI3-bound HH-responsive GBR with reduced acetylation in E9.25 WT limb buds compared to E10.5 WT limbs (note that the levels of enrichment are comparable to E10.5 *Shh^-/-^* limbs). B. Scatter plot of H3K27ac enrichment at HH-responsive GBRs in pre- and post-HH WT limbs. HH-responsive GBRs with significant reductions (FDR<0.05) in H3K27ac at E9.25 compared to E10.5, are denoted in dark blue, while the 3 GBRs with significantly higher H3K27ac in E9.25 limbs are in orange (FDR<0.05 n=2). C. Heatmap indicating relative H3K27ac enrichment at HH responsive GBRs in pre- and post-HH limbs (n=2). D. GLI3, HDAC1 and HDAC2 CUT&RUN tracks at E9.25 GLI-bound regions, pre- and post-HH. E. HDAC localization at E10.5 bound GBRs. Prior to HH signaling, most regions are enriched for both HDAC1 and HDAC2, while after, most GBRs are enriched for only HDAC1 (E9.25 HDAC1/2 n=2; E10.5 HDAC1/2 n=3). F. Initially most HDAC1-bound regions are also enriched for HDAC2, while after HH-signaling, fewer HDAC1-bound regions are also enriched for HDAC2. G. Prior to HH signaling, GLI3-bound regions are enriched for poised enhancer marks with lower levels of H3K27ac and HDAC enrichment. These are comparable to post-HH GLI3 enhancers where GLI3 actively represses targets. Orange shading in tracks indicates HH-responsive GBRs defined in Figure S1C. Scale bars =1kb. See also Figure S2, Supplemental File 1,2.

Previously, we found that GLI-mediated repression is facilitated by HDACs that regulate H3K27ac enrichment at GBRs (Lex et al. 2020). To determine whether GLI3 binding is associated with HDACs in pre-HH limb buds, we performed CUT&RUN in pre- and post-HH limbs with antibodies against HDAC1 and HDAC2. Pre-HH limb buds had similar levels of HDAC1 and HDAC2, with many GBRs enriched for both HDACs (Figure 2D,E). In post-HH limb buds, most of these regions continue to be bound by HDAC1 while HDAC2 binding is greatly reduced (Figure 2F). Overall, the presence of HDACs at most GBRs prior to HH is consistent with a model in which GLI3, together with an HDAC-containing repression complex, could be repressing enhancers to prevent premature activation of target genes (Figure 2G).

### GLI3 enhancers are unable to be prematurely activated with loss of *Gli3*

We next tested the hypothesis that the reduction in acetylation at GBRs prior to HH signaling was due to GLI3-mediated repression which reduces the levels of H3K27ac, preventing premature activation of enhancers (Figure 2G). We examined H3K27ac enrichment in WT and *Gli3^-/-^* pre-HH limb buds (Figure 3A), predicting that loss of GLI3-R at this time would result in increased acetylation at GBRs, as it does in E10.5 post-HH limbs (Lex et al., 2020) (Figure 3B). Contrary to our hypothesis, there was no significant increase in H3K27ac enrichment in *Gli3^-/-^* limb buds prior to HH induction (Figure 3B-F). Since GLI3-mediated repression of *Hand2* in the early limb is a key component of the pre-patterning model of anterior-posterior polarity (Litingtung et al., 2002; Osterwalder et al., 2014; Panman and Zeller, 2003; Welscher et al., 2002; Zhulyn et al., 2014; Bénazet and Zeller, 2009; Galli et al., 2010), we examined two GLI3 binding regions located 10kb and 85kb downstream of *Hand2* that have been suggested to mediate GLI3 repression (Vokes et al., 2008), but did not observe any increases in acetylation (Figure 3E, Figure S2A). We also considered the possibility that many enhancers, which have low levels of H3K27ac, may not be competent for activation at this stage and would therefore not have increased enrichment of H3K27ac upon loss of *Gli3*. To address this, we examined three outlier HH-responsive GBRs that contained significantly higher levels of H3K27ac in pre-HH limb buds (Figure 2B) to see if the acetylation levels might further increase in the absence of GLI3 repression. Despite GLI3 binding at these regions, there was no increase in H3K27ac enrichment with loss of *Gli3* before HH signaling (Figure S2B-D). Overall, this surprising result indicates that despite the presence of GLI3 and HDACs, GLI repression does not regulate H3K27ac enrichment at enhancers in pre-HH limb buds (Figure 3F).

**Figure 3.**
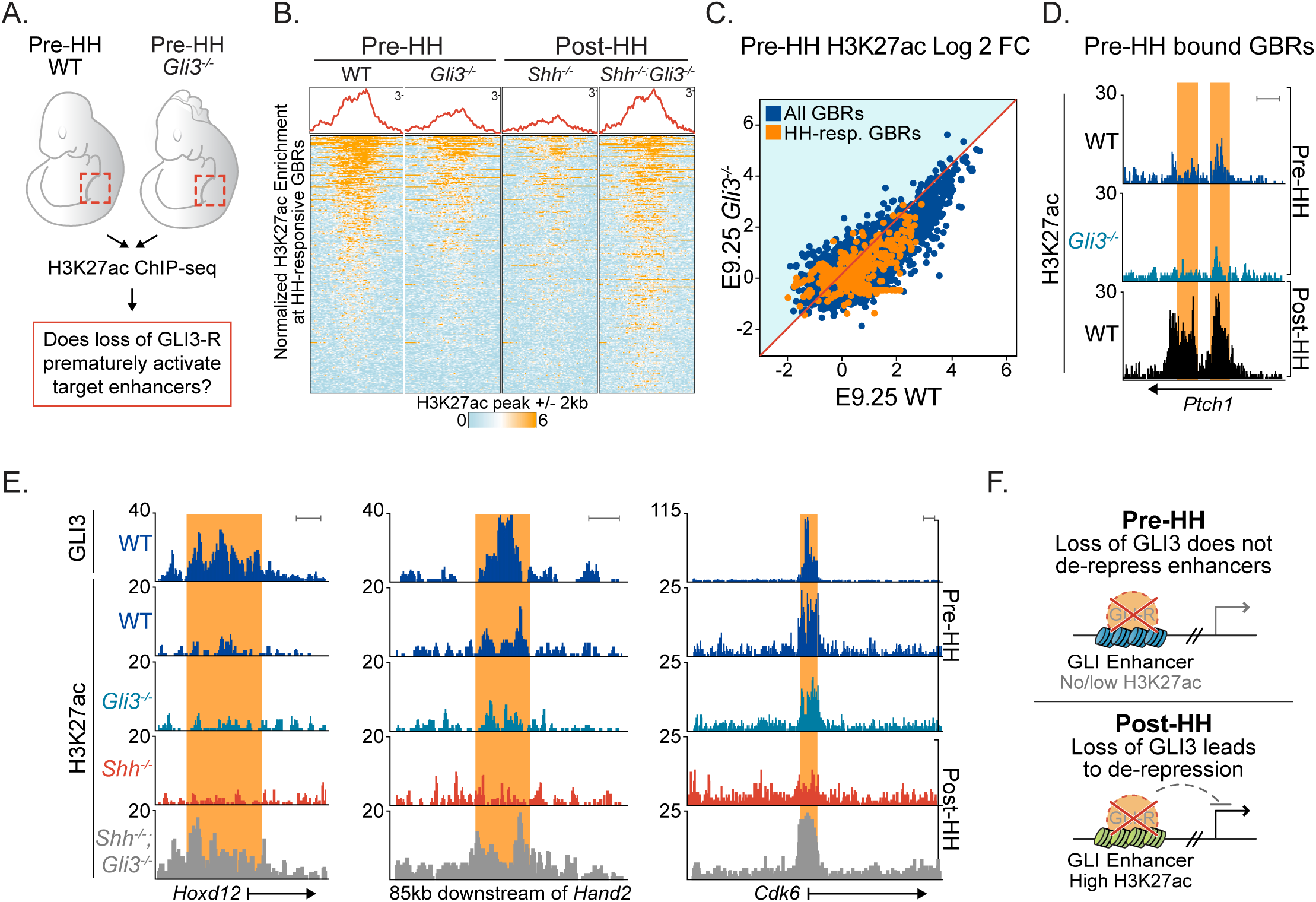
Loss of *Gli3* does not result in premature de-repression of enhancers. A. Schematic for testing whether loss of *Gli3* prematurely increases H3K27ac levels through GLI3 de-repression. B. Heatmap of H3K27ac enrichment at HH-responsive GBRs in pre-HH (E9.25, 21-23S) and post-HH (E10.5, 33-34S) limb buds, with loss of *Gli3*. At E10.5, *Shh^-/-^* limbs have reduced H3K27ac due to GLI3 repression. Loss of *Gli3* in *Shh^-/-^;Gli3^-/-^* results in de-repression of target enhancers and increased acetylation. This change is not observed prior to HH signaling with loss of *Gli3* compared to WT limb buds. C. Scatterplot of H3K27ac enrichment in pre-HH WT vs. *Gli3^-/-^* limbs shows no increase in acetylation with loss of *Gli3*. D. H3K27ac ChIP-seq tracks at a pre-HH GLI3-bound HH-responsive GBR with low H3K27ac in E9.25 WT limb buds that does not increase acetylation levels in E9.25 *Gli3^-/-^.* E. Examples of regions, bound by GLI3 at E9.25, that do not increase H3K27ac levels with loss of *Gli3* in pre-HH limb buds as they do in *Shh^-/-^;Gli3^-/-^* post-HH limbs. F. Schematic depicting loss of *Gli3* in pre-HH limb buds does not lead to de-repression of target enhancers as it does after HH induction at E10.5. Orange shading on tracks indicates HH-responsive GBRs defined in Figure S1C. Scale bars =1kb. See also Supplemental File 1,2.

### Most GLI3 target genes are unable to be prematurely activated with loss of *Gli3*

To determine whether the lack of enhancer activation in E9.25, pre-HH *Gli3^-/-^* limbs corresponded with a lack of de-repression of GLI target genes, we performed bulk RNA-seq on pre-HH WT and *Gli3^-/-^* limbs, as well as the anterior halves of post-HH (32-35S) WT and *Gli3^-/-^* limbs (Figure 4A-D). In post-HH *Gli3^-/-^* limbs, we identified a total of 159 significantly upregulated genes (FDR<0.05) that included well known signature HH target genes *Ptch1*, *Hand2*, *Hoxd13* and *Grem1* (Figure 4C,D) (Lewandowski et al., 2015). However, in agreement with the lack of GLI3-regulated enhancer activity, very few genes (9) were significantly upregulated with loss of *Gli3* in the early limb (Figure 4B). These genes include *Foxf1 and Osr1,* which are regulated by HH signaling in multiple mesodermal tissues but, with the exception of *Gpx6*, are not upregulated in E10.5 *Gli3^-/-^* limbs (Guzzetta et al., 2020; Hoffmann et al., 2014; Mahlapuu et al., 2001; Nasr et al., 2020). This suggests that rather than being a product of ongoing GLI3 repression, these upregulated genes may either represent contaminating flank tissue from the dissection. Alternatively, these could be residual mRNAs upregulated prior to limb bud initiation, as limb outgrowth requires cells from the lateral plate mesoderm to undergo an epithelial to mesenchymal transition which would have occurred within the last ∼8 hours (Gros and Tabin, 2014). Consistent with the former possibility, *Foxf1* is present in the lateral plate mesoderm but is not detected in either WT or *Gli3^-/-^* pre-HH limb buds (Figure 4D). Similarly, *Ptch1*, a signature HH target gene was detected in the neural tube, but was also not detected in the early limb buds in WT or *Gli3*^-/-^ embryos (Figure 4D). In support of the latter possibility, pre-HH upregulated mRNAs are reported to have an average half-life of 11.2 hours in differentiating ES cells (Figure S3A) (Sharova et al., 2009), and also have lower levels of retained introns than post-HH upregulated *Gli3^-/-^* genes, suggesting that some of the these transcripts may represent older mRNAs, potentially regulated by GLI3 in the lateral plate mesoderm, that are not currently being transcribed in the limb (Figure S3B,C). We conclude that GLI3 is unlikely to be engaged in ongoing transcriptional repression in the early limb bud.

**Figure 4.**
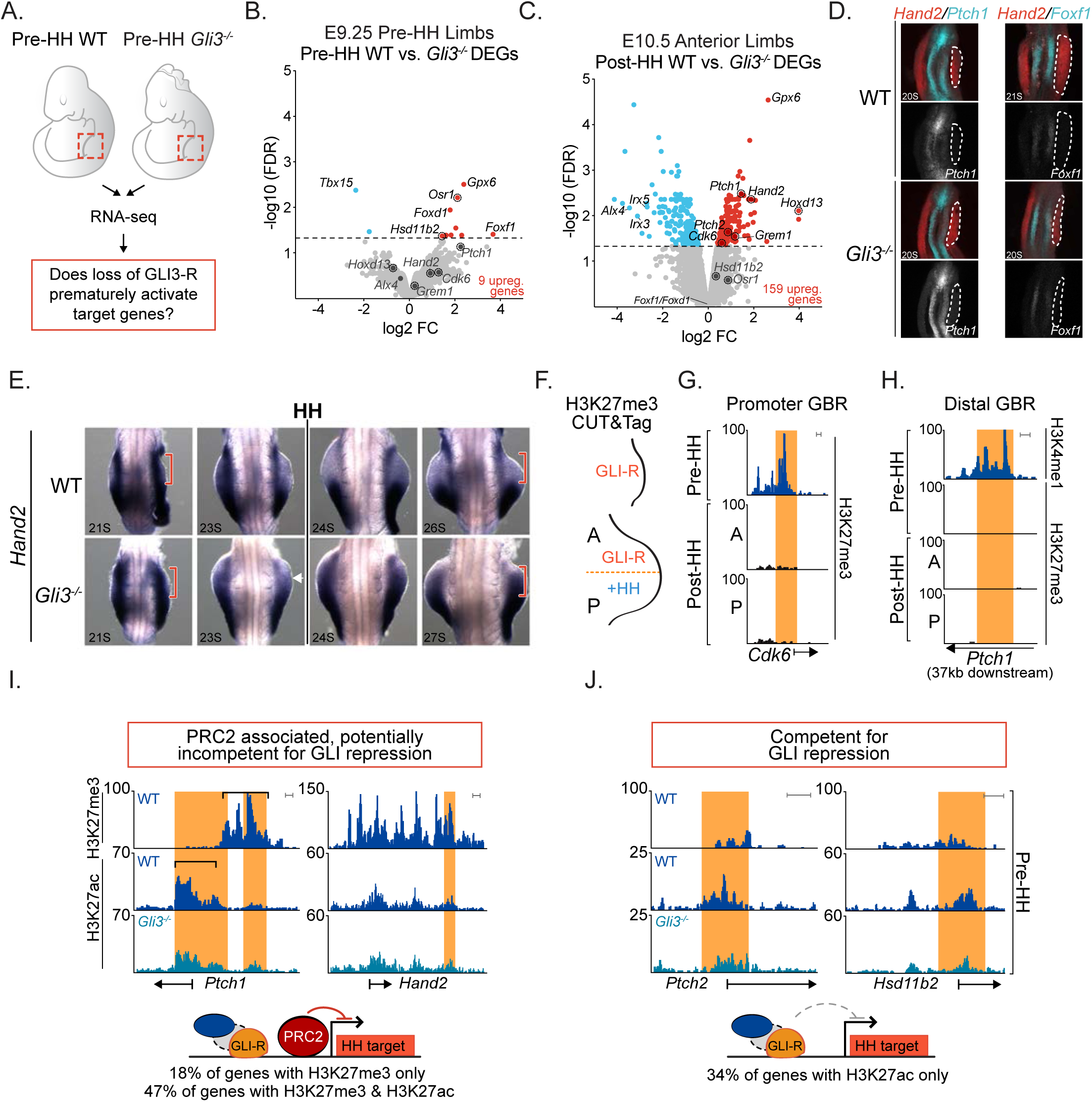
Most genes are not de-repressed in *Gli3* mutants prior to HH induction. A. Schematic for testing whether loss of *Gli3* can prematurely activate target genes through GLI3 de-repression. B,C. Volcano plot of differentially expressed genes (DEGs) detected in RNA-seq between WT and *Gli3^-/-^* limbs E9.25 (21-23S) and E10.5 (32-35S) anterior limb buds. D. Fluorescent in situ hybridization showing maximum intensity projections for *Hand2, Foxf1* and *Ptch1* in E9.25 WT and *Gli3^-/-^* limb buds indicating the absence of detectable *Foxf1 and Ptch1* in pre-HH limb buds. Dashed white lines outline the limb bud region. E. In situ hybridization for the GLI3 target *Hand2*, in WT and *Gli3^-/-^* limbs pre-(21, 23S) and immediately post-HH induction (24-27S; n=3). Note that *Hand2* is expressed almost uniformly across the limb bud in 21S embryos in both WT and *Gli3^-/-^* limbs. At 23S, slight reduction in anterior expression of *Hand2* is observed in both WT and *Gli3^-/-^* limbs (white arrowhead). The red brackets denote anterior limb. F. Schematic identifying H3K27me3 enriched regions in E9.25, E10.5 anterior (GLI3-repressed) and E10.5 posterior (HH signaling, loss of GLI-R) limb buds using CUT&Tag. G-J Orange shading in tracks indicates HH-responsive GBRs defined in Figure S1C. G. Example of a HH-responsive GBR at a HH target gene promoter with H3K27me3 enrichment specifically at E9.25 but not E10.5. H. Representative distal limb-specific HH-responsive GBR downstream of *Ptch1* with no H3K27me3 enrichment in E9.25 or E10.5 limb buds. I. Examples of HH target genes containing polycomb repression that could potentially be incompetent for GLI3 repression. J. Examples of genes that are bound by GLI3 at E9.25, contain H3K27ac enrichment and lack H3K27me3 enrichment, suggesting they are competent for GLI3 repression but are not de-repressed in the absence of *Gli3*. Scale bars for tracks indicate 1kb. See also Figure S3,S4, Supplemental File 2,3.

As *Hand2* is known to be repressed by GLI3 in early limb buds, we examined *Hand2* expression in pre-HH and early post-HH WT and *Gli3^-/-^* embryos to look for subtle changes that might not be detected by RNA-seq. *Hand2* is initially expressed throughout anterior WT limb buds (Figure 4D,E) similar to *Gli3^-/-^* limb buds (20-23S) (n=7). At early post-HH stages (24-26S), *Hand2* expression is variable in WT and *Gli3^-/-^* limb buds, with most WT limb buds expressing anterior *Hand2* and lower levels of anterior *Hand2* expression present even in some *Gli3^-/-^* limb buds (Figure 4E, white arrow). Posterior restriction of *Hand2* is not evident until 26S and is still variable at that time (Figure 4E, Figure S3D). Our findings are consistent with previous reports that noted early co-expression of *Gli3* and *Hand2* at these stages (Osterwalder et al., 2014). We conclude that GLI3 repression of *Hand2* is absent prior to the activation of HH signaling and suggests that GLI3 repression is first activated between 24-26S, coinciding with the initiation of HH signaling (Figure S1A,B).

### GLI3 repressor is inert in early limb buds

Our results thus far indicated that GLI repression is not active in the early limb bud. This could either be caused by inert GLI repression complexes or by target genes that were incapable of being activated. To understand whether PRC2 might broadly regulate HH targets, we computationally predicted 74 GLI target genes through identifying genes downregulated in *Shh^-/-^* limbs that were near HH-responsive GBRs (Supplemental Table 1)(Lewandowski et al., 2015). We performed CUT&Tag (Kaya-Okur et al., 2019) for H3K27me3, a modification indicative of PRC2 repression in pre-HH and post-HH limb buds (Figure 4F) (Margueron and Reinberg, 2011). There was little H3K27me3 enrichment at target gene promoters after HH induction, with the notable exceptions of *Ptch1* and *Gli1* in anterior limbs, consistent with previous reports (Lex et al., 2020) (Figure 4H,I; Figure S4A-D; Supplemental Table 1). However, in pre-HH limb buds, 81% of putative HH target gene promoters were enriched for H3K27me3 (Figure S4A). This H3K27me3 enrichment appears to resolve over limb development, as many genes completely lose H3K27me3 enrichment or have drastic reductions by E10.5, with only 21% remaining trimethylated in anterior limbs (Figure 4H; Figure S4A-D). Most HH-responsive GBRs lacked H3K27me3 enrichment, while the few GBRs enriched for H3K27me3, were generally located proximal to promoters (Figure 4I; Figure S4B). There were high levels of H3K27me3 at the *Hand2* locus in the early limb (Figure S4F), which as noted above is widely expressed throughout the limb at this time. *Hand2* also was enriched for H3K27ac, which is mutually exclusive with H3K27me3 (Zhang et al., 2015), suggesting that PRC2 is associated with *Hand2* in a subset of cells that presumably do not express *Hand2* in pre-HH limb buds (Figure S4F). Similar to *Hand2*, nearly half of predicted HH target genes were enriched for both H3K27ac and H3K27me3, which was also true of the few E10.5 genes enriched for H3K27me3 (Figure S4D; Supplemental Table 1). An important distinction between the two developmental stages is that at E9.25, most genes have high levels of H3K27me3 and low levels of H3K27ac, while at E10.5 when trimethylation is resolving, most genes have increased levels of H3K27ac with low levels of H3K27me3, consistent with increases in their expression levels at E10.5 (Figure S4D-G). For other genes enriched for both marks, like *Ptch1*, the distribution of these modifications is offset, supporting the potential for these mutually exclusive marks to be present at the same locus (Figure 4I, black brackets). The dual enrichment of H3K27ac and H3K27me3 may enable fast induction of these genes upon presence of relevant stimuli, while preventing inappropriate expression of them from non-relevant stimuli, as others have proposed (Kitazawa et al., 2021). A smaller population of genes (18%), are highly enriched for H3K27me3 and completely lack acetylation. Based on these findings, at least some HH target genes with H3K27me3 enrichment at this stage may not be competent for de-repression upon loss of *Gli3* (Figure 4I) and GLI3 repression itself is unlikely to facilitate H3K27me3 enrichment (see discussion). Importantly, a third of genes are bound by GLI3 with no H3K27me3 enrichment and are likely to be competent for de-repression upon the loss of *Gli3*. However, as we did not observe increased acetylation in E9.25 *Gli3^-/-^* limbs at these regions as we do in E10.5 *Gli3^-/-^* limb buds, we conclude that GLI3 repressor is inert in the early limb bud (Figure 4J).

### GLI3-dependent chromatin compaction does not initiate until after the start of HH signaling

To define the onset of GLI repression, we looked for evidence of GLI3 repression at 28-30S (E10), about 8-12 hours after HH activation. We used chromatin accessibility as a read out for GLI repression, as E10.5 (35S) *Shh^-/-^* limb buds (constitutive GLI repression) have reduced accessibility at HH-responsive GBRs compared to WT limb buds (Lex et al., 2020). To verify that this chromatin compaction was mediated through GLI repression and not lack of GLI activator, we performed ATAC-seq in *Shh^-/-^;Gli3^-/-^* E10.5 limb buds. Loss of *Gli3* resulted in accessible chromatin at HH-responsive GBRs similar to WT controls, confirming that chromatin compaction is GLI3 repressor-dependent (Figure 5A).

**Figure 5.**
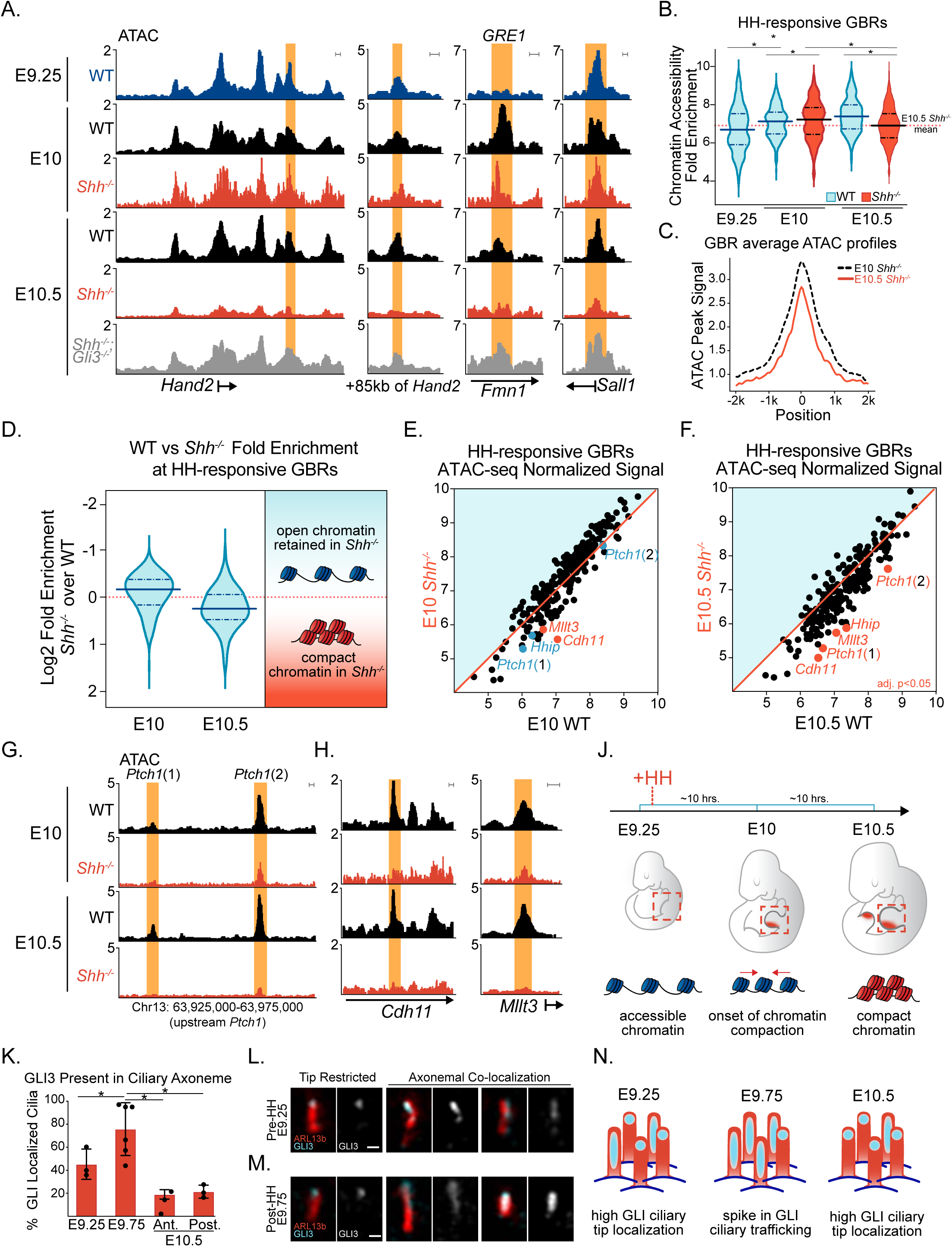
GLI3-dependent chromatin compaction occurs after HH induction. A. ATAC-seq tracks showing examples of HH-responsive GBRs that are compacted in E10.5 *Shh^-/-^* limbs but not in E9.25 WT or E10 *Shh^-/-^* limbs. Note E10.5 *Shh^-/-^;Gli3^-/-^* limbs maintain accessible chromatin at these regions. B. Violin plots of chromatin accessibility in WT and *Shh^-/-^* limbs. Globally the chromatin at HH-responsive GBRs is more accessible in E10 Shh^-/-^ limb buds compared to E10 WT limbs and E10.5 *Shh^-/-^* limbs (n=2-3 at each stage). Wilcoxon signed rank tests were performed to compare each pair of groups, multiple hypothesis testing adjusted using BH method. E9.25<E10 WT, FDR<2.69E-15; E9.25<E10 *Shh^-/-^* FDR<3.44E-23; E9.25<E10.5 FDR<5.82E-15 E10 WT<E10 *Shh^-/-,^* FDR<2.55E-4; E10 *Shh^-/-^>*E10.5 *Shh^-/-^*, FDR<1.21-18; E10.5 WT>E10.5 *Shh^-/-^,* FDR<9.81E-12). C. Average ATAC profiles of HH-responsive GBRs in E10 and E10.5 *Shh^-/-^* limb buds. D. Violin plots depicting log2 fold changes in chromatin accessibility (WT versus *Shh^-/-^*) at E10 and E10.5 E (p<0.0001 Wilcoxon signed rank test). E,F. Scatter plots of ATAC-seq signal in WT vs. *Shh^-/-^* limbs at E10 (E) and E10.5 (F). GBRs and their associated genes annotated in red signify a significant reduction of chromatin accessibility in *Shh^-/-^* limbs compared to WT (FDR<0.05). GBRs in blue indicate visibly reduced accessibility in E10 *Shh^-/-^* limb buds that are not significantly changed until E10.5 G. ATAC-seq tracks of GBRs near *Ptch1* that are reduced, but not significant, in E10 *Shh^-/-^* limbs compared to E10 WT limb buds. H. GBRs that are already have significantly reduced accessibility in E10 *Shh^-/-^* limbs compared to E10 WT. J. Schematic showing the temporal onset of chromatin compaction at GBRs in relation to the initiation of HH signaling. K. Quantification of GLI3 present along ciliary axonemes out of total number of cilia (marked by ARL13b) that colocalize with GLI3 in E9.25 (21-23S) (n=3), E9.75 (26-28S) center limb buds (n=6) and E10.5 (35S) anterior and posterior limbs (n=3). Error bars indicate SEM. L. Representative images of GLI3^FLAG^ ciliary distribution in E9.25 and E9.75 limb buds acquired on a Zeiss LSM 710 Confocal visualized using maximal intensity projections. N. Schematic depicting the temporal shift in GLI3 ciliary localization. Orange shading in all tracks indicates HH-responsive GBRs defined in Figure S1C. Scale bars for ATAC-seq tracks =1 kb. Scale bars for panel L =0.5μm. See also Figure S5, Supplemental File 4.

As *Hand2* expression becomes posteriorly restricted in the limb around 26S, we anticipated that GLI3 repression should be established by 28-30S (E10), and predicted that chromatin in *Shh^-/-^* limbs at this stage would be more compacted compared to WT. Alternatively, if GLI-repression was yet to become established, or was still in the process of being established, the accessibility of HH-responsive GBRs would be similar to that of E10 WT limbs. Consistent with the latter scenario, the chromatin at HH-responsive GBRs overall was not compacted. In fact, it was significantly more accessible in E10 *Shh*^-/-^ limb buds compared with E10 WT limbs (Figure 5A-D). This surprising result indicated that one of the signatures of GLI repression had largely not been established in this population at this stage in development. We next examined individual GBRs to determine if there were any regions with less accessible chromatin in E10 *Shh*^-/-^ limb buds, indicative of initial targets of GLI3 repression. While most regions were not significantly reduced, several regions have a trend toward having reduced accessibility at E10 in *Shh^-/-^* compared to WT, such as HH-responsive GBRs near HH targets *Ptch1* and *Hhip*. However, this reduction did not become significant until E10.5 (Figure 5E-G, blue datapoints). Only a few HH-responsive GBRs had significantly reduced accessibility in E10 in *Shh^-/-^* limb buds including GBRs around *Cdh11* and *Mllt3*, which were also significantly reduced at E10.5 (Figure 5E,F,H; Supplemental File 4). Collectively these results support that GLI repression is not fully established even up to 12 hours after HH signaling would have normally become active (Figure 5J).

### Changes in GLI3 ciliary distribution correspond with the onset of GLI3-dependent chromatin compaction

As GLI3 repression does not initiate until after the induction of HH signaling, we asked if there was a change in the ciliary trafficking of GLI3 proteins that could signify changes in the processing of GLI3. In pre-HH limb buds, ∼80% of cilia contained endogenous GLI3^FLAG^ (Figure S5A,B,E). There were similarly high trends in both GLI3, as well as GLI2 ciliary co-localization in pre-HH WT and *Shh^-/-^* limb buds, where expression was primarily localized to the ciliary tip (Fig. S5A-E). This indicates that this enrichment is not due to undetectable levels of SHH that might still promote GLI activation and, as GLI-R is not localized to the cilia, it suggests that pre-HH ciliary localization is enriched with full-length, unprocessed GLI protein (Figure S5A-E) (Haycraft et al., 2005; Wen et al., 2010). While GLI ciliary localization remained at comparable levels shortly after the onset of SHH expression (E9.75, 26-28S) as well as at E10.5 (32-35S; Fig. S5F), E9.75 limb buds had a significant redistribution of GLI3 within the cilia, with 74% of cilia having GLI3 signal along the ciliary axoneme and a corresponding reduction in the percentage of cilia with tip-restricted GLI3 expression (Fig. 5K, L; Fig. S5G). Overall, this data is consistent with the possibility that there may be a ‘steady state’ of low to moderate levels of GLI3 ciliary trafficking and processing, with a spike in the rate of trafficking coinciding with the requirement for GLI3 repression to become established during limb development.

### GLI3 is unlikely to co-regulate HAND2 targets in early limb buds despite co-expression

GLI3-repression of *Hand2* is initiated concomitantly with activation of HH signaling and therefore GLI3 does not have a bona-fide pre-patterning role. In pre- and post-HH limb buds, GLI3 and HAND2 bind to nearly adjacent regions in IRX3/5 as well as many other targets (Osterwalder, 2014; Vokes et al, 2008), suggesting that they might co-regulate a common set of binding regions. (Figure S6A) (Osterwalder et al., 2014). Interestingly, HAND2 and GLI3 have been shown to bind to each other and to synergistically activate targets through common CRMs in developing facial elements (Elliott et al., 2020). While HAND2 and GLI3 are mutually exclusive in later limb buds (teWelscher et al, 2002; Osterwalder et al, 2014), their co-occurrence in pre-HH limb buds suggests the possibility that GLI3-R could facilitate HAND2 regulation of target genes in its capacity as a DNA binding protein rather than as a repressor. In support of this possibility, HAND2-bound enhancers for *Tbx2* and *Tbx3* contain GBRs that have called GLI3 peaks in pre-HH but not post-HH limb buds (Figure S6B, light blue shading). To determine whether GLI3 and HAND2 might initially co-activate targets in the early limb in a similar fashion to craniofacial tissue, we identified a population of 310 competent GBRs specifically bound by GLI3 at E9.25, but not at E10.5, that also overlap previously identified HAND2 limb binding sites (Figure S6B,C)(Osterwalder et al., 2014). We analyzed previously reported HAND2 and GLI3 motifs (Elliott et al., 2020; Lex et al., 2020) in ATAC-seq footprints in the early limb compared to post-HH *Shh^-/-^* limbs, which have active GLI repression and would not be expected to have HAND2 and GLI3 co-regulation. If GLI3 and HAND2 were to coordinate enhancer activation, we predicted HAND2 motifs would be enriched in pre-HH limbs compared to post-HH *Shh^-/-^* limbs. Inconsistent with this scenario, there was no difference in HAND2 motif enrichment between pre-HH WT and post-HH *Shh^-/-^* limbs (Figure S6C-E, Fisher’s extract test p-value= 0.14). Since *Hand2* is only required for limb bud initiation (Galli et al., 2010), the absence of increased motif enrichment in pre-HH limb buds does not support a role for HAND2 in regulating this population of GLI3-bound early enhancers. We also note that *Tbx2* and *Tbx3* are not reduced in pre-HH *Gli3^-/-^* limb buds as would be expected if GLI3 had a significant role in co-activating these HAND2 target genes (Supplemental File 3).

## Discussion

GLI repressors have been proposed to play a significant role in ‘pre-patterning’ the anterior-posterior limb bud prior to the onset of HH signaling (Chiang et al., 2001; Litingtung et al., 2002; Osterwalder et al., 2014; Panman and Zeller, 2003; te Welscher et al., 2002; Zhulyn et al., 2014; Wyngaarden et al., 2011; reviewed in Zuniga and Zeller, 2020). Unexpectedly, we find that GLI3 does not act as a transcriptional repressor in the early limb bud before the onset of HH signaling. Although the GLI3-R isoform is produced at comparable levels and binds to chromatin, it does not mediate deacetylation of H3K27ac or chromatin compaction at enhancers, as it does after HH signaling (Lex et al., 2020). Moreover, there is little to no upregulated gene expression in early *Gli3^-/-^* limb buds. The lack of GLI repressor activity prior to the onset of Hedgehog signaling is inconsistent with the current pre-patterning model, instead it suggests that the repressive patterning activities of GLI3 in the limb bud are initiated concurrently with, or after, the initiation of HH signaling (Figure 6). This suggests that the widespread alterations in gene expression present in ciliopathies are primarily caused by misexpression of normally repressed GLI3 target genes during ongoing Hedgehog signaling rather than by precocious de-repression.

**Figure 6.**
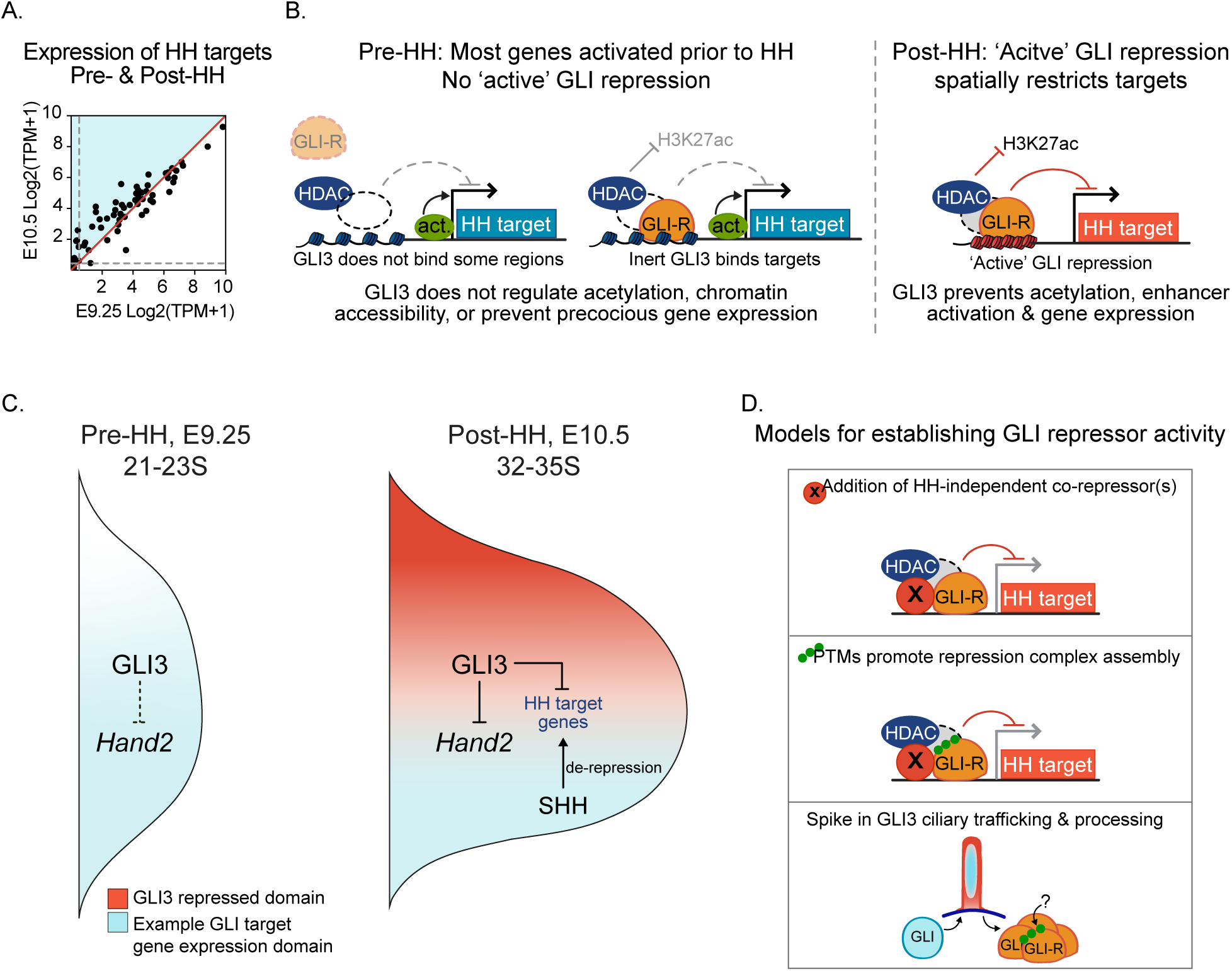
Model for established GLI3-mediated repression. A. Comparison of expression levels of predicted GLI target genes (Supplemental File 3), with pre-HH H3K27ac enrichment at promoters (60/74 genes) in pre-(21-23S) and post-HH (32-35S) limb buds. Grey dashed lines are indicative of minimal expression thresholds, log2(TPM+1)<0.5. Note that most GLI target genes are already transcribed in pre-HH limb buds. B. Model for lack of GLI3 repression prior to HH signaling. In pre-HH limb buds, many target genes are already activated by HH-independent factors. While GLI3 binds to many (but not all) targets, it appears inert as it is unable to regulate deposition of H3K27ac at enhancers or chromatin accessibility as it does in later limb development. After HH induction when ‘active’ GLI3 repression has been established, GLI3 prevents addition of H3K27ac at enhancers to spatially restrict expression of its target genes and pattern the limb. C. Schematic of GLI3 spatial regulation of target genes. In pre-HH limb buds, GLI3 does not repress targets such as *Hand2*, which is expressed in an expanded anterior domain throughout the limb bud. In contrast, GLI3 restricts expression of targets to the posterior limb bud where HH signaling is active in post-HH limb buds. D. Possible models for initiating GLI3 repressor activity. Top: HH-independent co-repressor(s), which may not be abundant in early limb development collaborate with GLI3 to assemble a complete repression complex. Middle: Addition of post-translational modifications (PTMs), potentially added via ciliary trafficking and processing, promote assembly of, or stabilize, a GLI repression complex. Bottom: A spike in ciliary GLI3 processing, as reflected by increased axonemal colocalization, increases the amount of viable GLI3 repressor, potentially through addition of post-translation modifications as suggested above.

### GLI3-R is transcriptionally inert in early limb buds

The absence of GLI repression in the early limb bud could be caused by inert GLI-R isoforms or by target genes that are not yet competent for GLI repression (Figure 4I,J). Our results are consistent with the co-occurrence of both of these mechanisms. GLI3 does not bind to almost a third of all HH-responsive GBRs that will be bound at E10.5, and most unbound regions having not yet acquired a poised conformation (Figure 1; Figure S1G,H). In this case, pioneer factors may be required to prime this subset of GBRs before GLI3 can regulate them, as has been proposed for SOX2’s role in activating GLI neural enhancers (Oosterveen et al., 2012; Peterson et al., 2012). The presence of the polycomb repressive mark, H3K27me3, at the promoters of many predicted HH target genes, provides an additional mechanism of GLI3-independent repression, as it is highly enriched at many promoters in E9.25 limbs, but is greatly reduced or absent from post-HH limb buds (Figures 4F-J, S4). Notably, GLI3 de-repression is unlikely to mediate this reduction, as many of these genes also lack H3K27me3 in E10.5 *Shh^-/-^* limb buds (Lex et al., 2020). This suggests that removal of H3K27me3 is largely independent of HH signaling and GLI3 regulation, despite reports demonstrating that H3K27me3 enrichment at some HH targets is resolved in a HH-dependent manner (Figures 4F-J, S4)(Shi et al., 2014). The H3K27me3 clearing at these genes is most likely mediated by the loss of unknown, GLI-independent transcriptional repressors as has been proposed within the developing brain (He et al., 2020).

Unlike the previous examples, the class of genes that are bound by GLI3 and not enriched for H3K27me3 are more likely to be competent for GLI repression in pre-HH limb buds (Figure 4J). However, these genes are also not upregulated in *Gli3^-/-^* limb buds (Fig. 4B,D), suggesting an absence of GLI3 repressor function at this stage. Several additional lines of evidence support this suggestion. First, *Hand2* is broadly expressed in the anterior domain of pre-HH limb buds where there is co-localization with GLI3-R that persists through E9.5 (Figure 4D,E; Osterwalder et al., 2014). Second, H3K27ac levels at HH responsive GBRs do not increase in pre-HH *Gli3^-/-^* limb buds (Figure 3) as they do in post-HH *Gli3^-/-^* limb buds, even at genes that appear competent for GLI-repression (Figure 3B,E; 4J; Lex et al., 2020). Third, while most HH-responsive enhancers have lower H3K27ac enrichment in pre-HH limb buds than in post-HH limb buds (Figure 2B,C), exceptional enhancers with higher levels of H3K27ac do not further increase acetylation levels in *Gli3^-/-^* limb buds, excluding the possibility that lack of increased H3K27ac levels are due to lack of a HH-independent activator (Figure S2B-D). Finally, HH-responsive GBRs are broadly accessible at pre-HH stages and up until ∼10 hours after the time of HH induction, in contrast to the loss of accessibility seen under conditions of maximal GLI repression in post-HH E10.5 limbs (Figure 5).

### Mechanisms for establishing GLI-repressor activity

One mechanism for regulating repressor activity is the variable expression of required co-factors as exemplified by the Brinker repressor (Upadhyai and Campbell, 2013). Consistent with this scenario, the genes encoding several co-repressors implicated with GLI repression in various contexts, including *Ski*, *Smarcc1* and *Atrophin1* (Dai et al., 2002; Jeon and Seong, 2016; Zhang et al., 2013) are significantly downregulated prior to HH induction compared to at E10.5 (Supplemental File 3). Conversely, a protein enriched in early limb buds could inhibit GLI3 repressor activity, as has been shown for HOXD12 in post-HH limb buds (Chen et al., 2004). Although HDACs are enriched at GBRs in both pre- and post-HH limb buds (Figure 2E,G), the lack of increased H3K27ac enrichment in *Gli3^-/-^* limb buds suggests that they are either inactive or not regulated by GLI3. Interestingly, both HDAC1 and HDAC2 initially co-localize at many GBRs at E9.25, while in post-HH limb buds, most remain bound by HDAC1 but lose HDAC2 enrichment. In other contexts, HDAC2 is required to recruit, but not maintain HDAC1 at some enhancers (Somanath et al. 2017), and a similar mechanism could enable the assembly of an HDAC1-containing GLI repression complex. As HDACs require co-repressor complexes to guide them to their substrates (reviewed in Adams et al., 2018), the absence of a functional GLI3 co-repressor might prevent them from being regulated by HDACs. Alternatively, rather than missing a co-repressor, it is possible that GLI3-R lacks additional, uncharacterized post-translational modifications required for its activity, or to functionally interact with its repression complex (Figure 6D).

### Temporal onset of GLI transcriptional repression

GLI-dependent restriction of chromatin accessibility can first be detected by ATAC-seq at 28-30S when a few HH-responsive GBRs have significantly reduced accessibility in *Shh^-/-^* limb buds. However, most regions are actually more accessible than in E10.5 *Shh^-/-^*, suggesting that GLI repression is not widespread at this time (Figure 5). Focusing on *Hand2* as a likely direct target of GLI3 repression (Osterwalder et al., 2014; Vokes et al., 2008; te Welscher et al., 2002), there is no reduction in chromatin accessibility at this locus in E10 *Shh^-/-^* limbs as is seen at E10.5 (Figure 5A). *Hand2* expression does not become posteriorly restricted until around 26S (Figure 4E), an observation that is consistent with previous findings (Osterwalder et al., 2014), suggesting that GLI3 transcriptional repression of *Hand2* commences around 26S, a timepoint coinciding with the activation of *Shh* expression in the limb bud (Figure 1A,B, Figure S1 A,B) (Charite et al., 2000; Zhu et al., 2008). The lack of detectable reductions in chromatin accessibility at a time shortly after repression of *Hand2* has initiated, indicates that GLI-dependent compaction likely occurs as a later step in GLI3-mediated repression.

The onset of GLI transcriptional repression is accompanied by a re-distribution of ciliary GLI3 into the axoneme, suggestive of a possible increase in GLI3 trafficking that coincides with the time when GLI3 starts to functionally repress target genes. We propose that increased GLI3 trafficking is facilitated by developmental changes to unknown ciliary processing components that regulate GLI3-R activity (Figure 6D). Since there are comparable levels of GLI3-R protein present within the pre- and post-HH limb buds (Fig. 1C), this regulation seems most likely to affect an unknown post-translational modification to GLI3-R rather than the processing of GLI3-FL into truncated GLI3-R (Figure 6D). If this is the case, then the activation of GLI repression is dependent on the status of the cilia and GLI transcriptional repression may be dynamically regulated in different developmental contexts.

### A Model for establishing GLI3-mediated repression

Our findings provide genetic and genomic evidence that GLI3-R is inert prior to HH induction in the limb bud. As GLI3-dependent chromatin compaction is established in E10.5, but not E10 *Shh^-/-^* limb buds, it also suggests that GLI repression is likely established temporally during limb development through HH-independent mechanisms. Rather than preventing initial expression of HH targets, most genes repressed by GLI3 in later limb development are initially expressed in pre-HH limb buds (Figure 6A; Supplemental File 3). Once repression is later established through unknown mechanisms, GLI3 functions to restrict target gene expression to the posterior limb (Figure 6B-D). More broadly, this work indicates that GLI repression is not a default state and instead must be acquired in context-dependent fashion. This has major implications for our understanding of the pathway itself and whether genes and enhancers must gain competency to be regulated by HH signaling.

## Acknowledgements

We thank Matt Anderson and Mark Lewandoski for sharing their protocol for clearing embryos, We thank the Henikoff Lab and EpiCypher for providing CUT&Tag reagents, and Jonathan Eggenschwiler for providing the GLI2 antibody. We thank Jessica Podnar from the Genomic Sequencing and Analysis Facility at the University of Texas at Austin, Anna Webb from the Center for Biomedical Research Support at the University of Texas at Austin for technical advice, and Janani Ramachandran for comments on this manuscript. This work was supported by NIH R01HD073151 (to SAV and HJ), F31DE027597 (to RKL) and R01HG009518 (to HJ).

## Author Contributions

Conceptualization, R.K.L. and S.A.V.; Methodology, R.K.L, W.Z., Z.J., Software J.D.K.; Validation K.E.W.; Formal Analysis, R.K.L., W.Z., Z.J.; Investigation, R.K.L., W.Z., Z.J., K.N.F., K.E.S., K.E.W.; Data curation, R.K.L., W.Z.; Writing – Original Draft, R.K.L. and S.A.V.; Writing – Review & Editing, S.A.V, R.K.L, W.Z., H.J.; Supervision, S.A.V., H.J.; Funding Acquisition, S.A.V., H.J., R.K.L.

## STAR Methods

### KEY RESOURCES TABLE

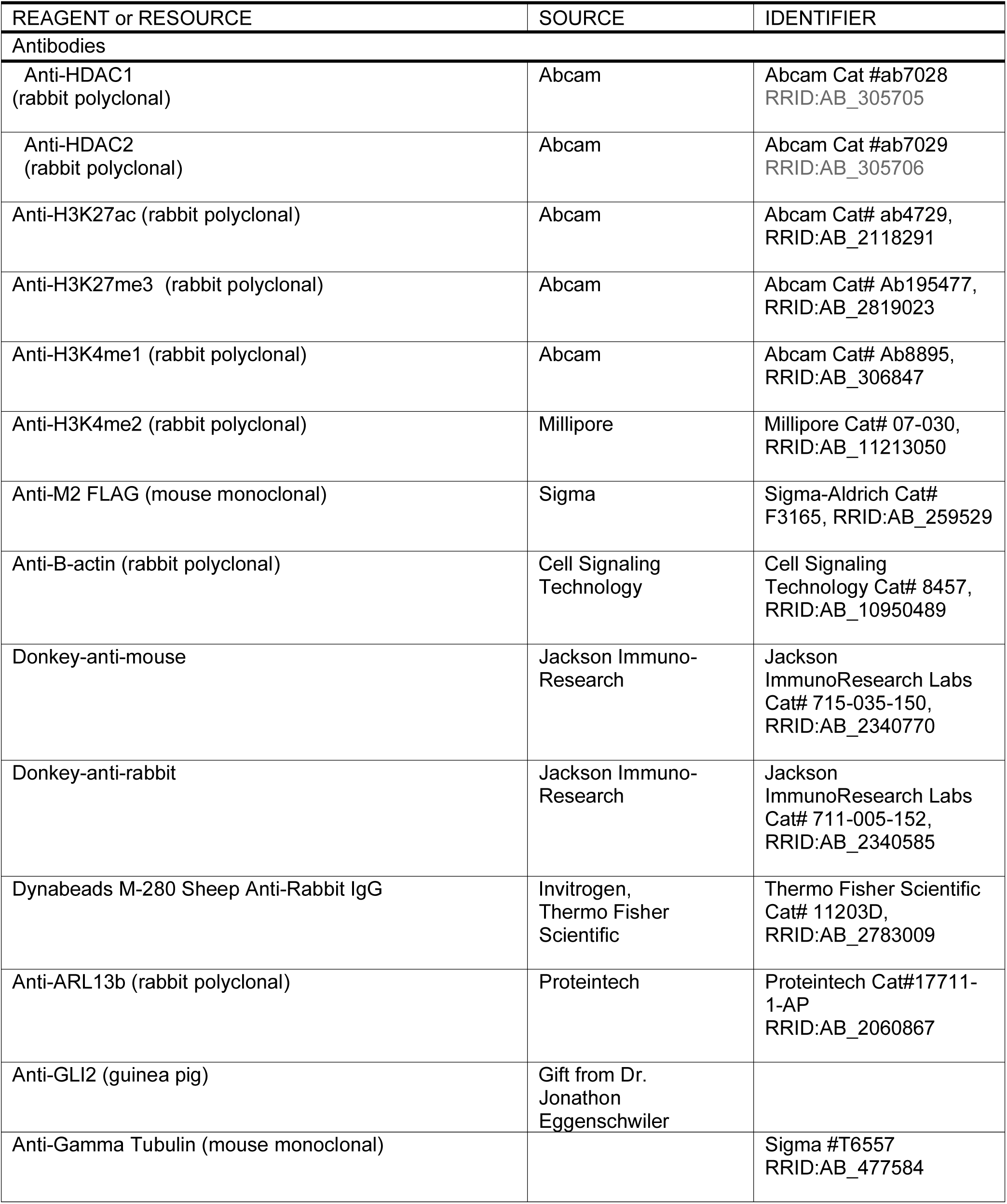
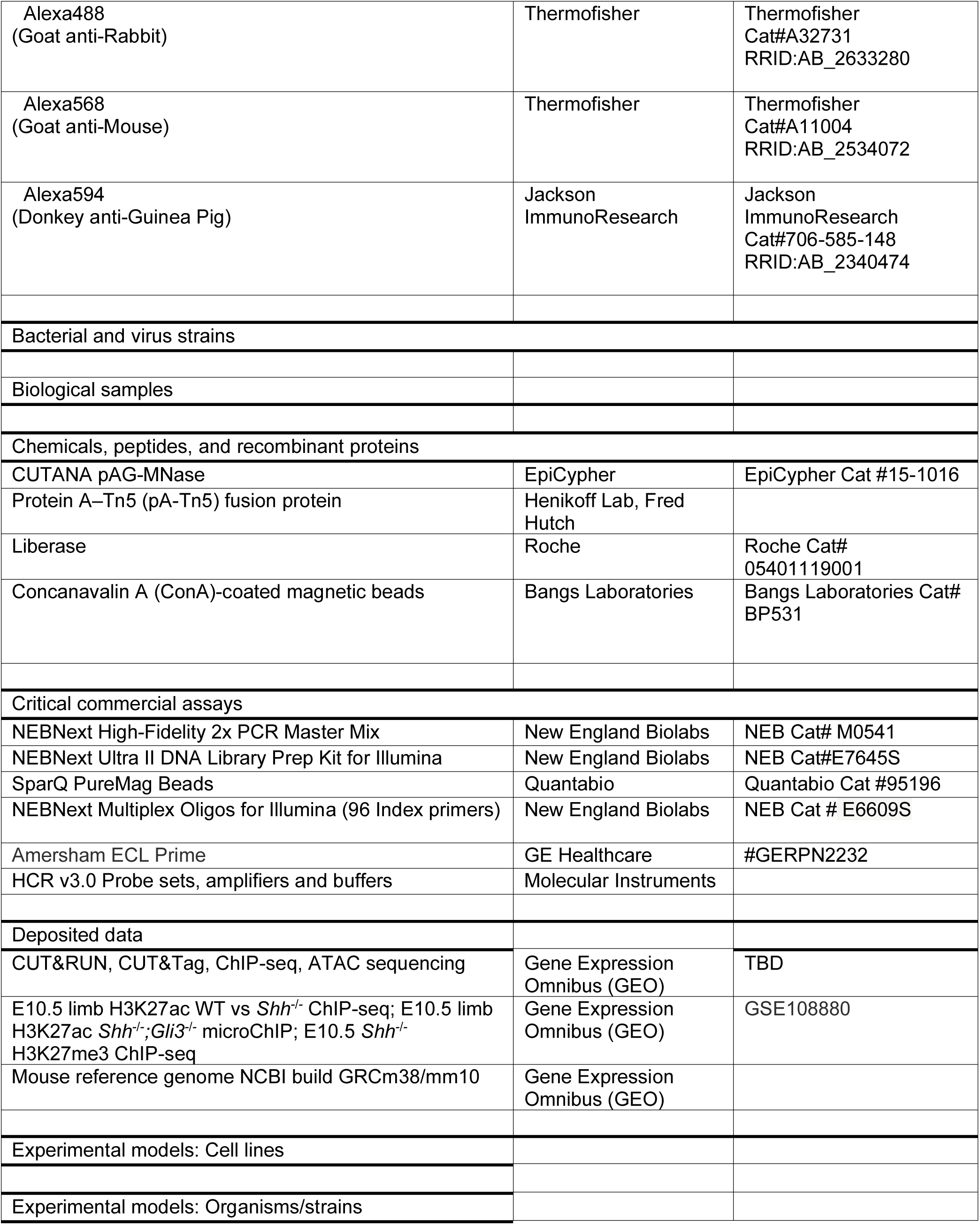
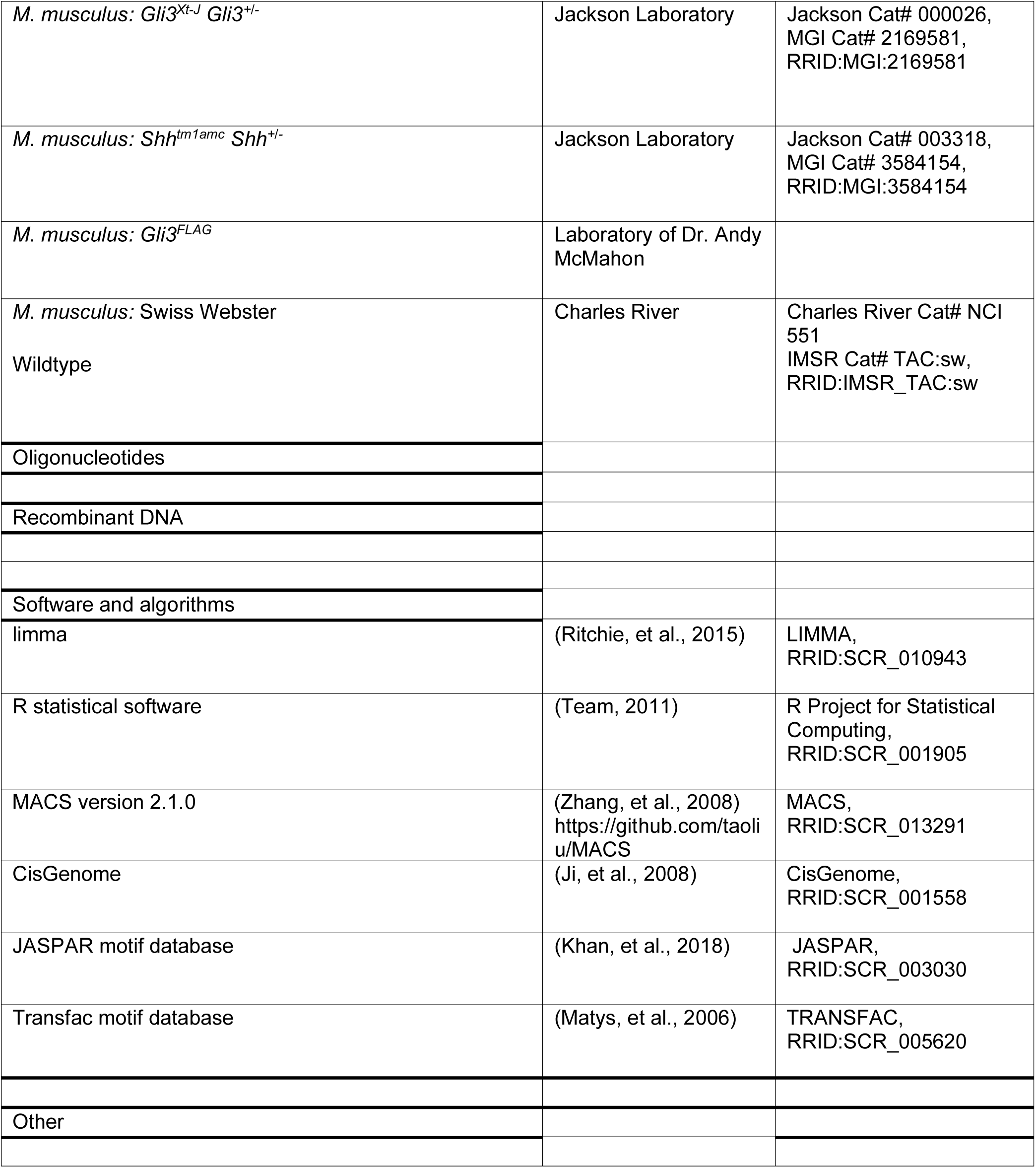

| REAGENT or RESOURCE | SOURCE | IDENTIFIER |
| --- | --- | --- |
| **Antibodies** |  |  |
| Anti-HDAC1<br>(rabbit polyclonal) | Abcam | Abcam Cat #ab7028<br>RRID:AB_305705 |
| Anti-HDAC2<br>(rabbit polyclonal) | Abcam | Abcam Cat #ab7029<br>RRID:AB_305706 |
| Anti-H3K27ac (rabbit polyclonal) | Abcam | Abcam Cat# ab4729,<br>RRID:AB_2118291 |
| Anti-H3K27me3 (rabbit polyclonal) | Abcam | Abcam Cat# Ab195477,<br>RRID:AB_2819023 |
| Anti-H3K4me1 (rabbit polyclonal) | Abcam | Abcam Cat# Ab8895,<br>RRID:AB_306847 |
| Anti-H3K4me2 (rabbit polyclonal) | Millipore | Millipore Cat# 07-030,<br>RRID:AB_11213050 |
| Anti-M2 FLAG (mouse monoclonal) | Sigma | Sigma-Aldrich Cat#<br>F3165, RRID:AB_259529 |
| Anti-B-actin (rabbit polyclonal) | Cell Signaling<br>Technology | Cell Signaling<br>Technology Cat# 8457,<br>RRID:AB_10950489 |
| Donkey-anti-mouse | Jackson Immuno-Research | Jackson<br>ImmunoResearch Labs<br>Cat# 715-035-150,<br>RRID:AB_2340770 |
| Donkey-anti-rabbit | Jackson Immuno-Research | Jackson<br>ImmunoResearch Labs<br>Cat# 711-005-152,<br>RRID:AB_2340585 |
| Dynabeads M-280 Sheep Anti-Rabbit IgG | Invitrogen,<br>Thermo Fisher<br>Scientific | Thermo Fisher Scientific<br>Cat# 11203D,<br>RRID:AB_2783009 |
| Anti-ARL13b (rabbit polyclonal) | Proteintech | Proteintech Cat#17711-1-AP<br>RRID:AB_2060867 |
| Anti-GLI2 (guinea pig) | Gift from Dr.<br>Jonathon<br>Eggenschwiler |  |
| Anti-Gamma Tubulin (mouse monoclonal) |  | Sigma #T6557<br>RRID:AB_477584 |
| Alexa488<br>(Goat anti-Rabbit) | Thermofisher | Thermofisher<br>Cat#A32731<br>RRID:AB_2633280 |
| Alexa568<br>(Goat anti-Mouse) | Thermofisher | Thermofisher<br>Cat#A11004<br>RRID:AB_2534072 |
| Alexa594<br>(Donkey anti-Guinea Pig) | Jackson<br>ImmunoResearch | Jackson<br>ImmunoResearch<br>Cat#706-585-148<br>RRID:AB_2340474 |
| Bacterial and virus strains |  |  |
| Biological samples |  |  |
| Chemicals, peptides, and recombinant proteins |  |  |
| CUTANA pAG-MNase | EpiCypher | EpiCypher Cat #15-1016 |
| Protein A-Tn5 (pA-Tn5) fusion protein | Henikoff Lab, Fred<br>Hutch |  |
| Liberase | Roche | Roche Cat#<br>05401119001 |
| Concanavalin A (ConA)-coated magnetic beads | Bangs Laboratories | Bangs Laboratories Cat#<br>BP531 |
| Critical commercial assays |  |  |
| NEBNext High-Fidelity 2x PCR Master Mix | New England Biolabs | NEB Cat# M0541 |
| NEBNext Ultra II DNA Library Prep Kit for Illumina | New England Biolabs | NEB Cat#E7645S |
| SparQ PureMag Beads | Quantabio | Quantabio Cat #95196 |
| NEBNext Multiplex Oligos for Illumina (96 Index primers) | New England Biolabs | NEB Cat # E6609S |
| Amersham ECL Prime | GE Healthcare | #GERPN2232 |
| HCR v3.0 Probe sets, amplifiers and buffers | Molecular Instruments |  |
| Deposited data |  |  |
| CUT&RUN, CUT&Tag, ChIP-seq, ATAC sequencing | Gene Expression<br>Omnibus (GEO) | TBD |
| E10.5 limb H3K27ac WT vs *Shh*<sup>-/-</sup> ChIP-seq; E10.5 limb<br>H3K27ac *Shh*<sup>-/-</sup>; *Gli3*<sup>-/-</sup> microChIP; E10.5 *Shh*<sup>-/-</sup><br>H3K27me3 ChIP-seq | Gene Expression<br>Omnibus (GEO) | GSE108880 |
| Mouse reference genome NCBI build GRCm38/mm10 | Gene Expression<br>Omnibus (GEO) |  |
| Experimental models: Cell lines |  |  |
| Experimental models: Organisms/strains |  |  |
| *M. musculus: Gli3<sup>Xt-J</sup> Gli3<sup>+/-</sup>* | Jackson Laboratory | Jackson Cat# 000026,<br>MGI Cat# 2169581,<br>RRID:MGI:2169581 |
| *M. musculus: Shh<sup>tm1amc</sup> Shh<sup>+/-</sup>* | Jackson Laboratory | Jackson Cat# 003318,<br>MGI Cat# 3584154,<br>RRID:MGI:3584154 |
| *M. musculus: Gli3<sup>FLAG</sup>* | Laboratory of Dr. Andy McMahon |  |
| *M. musculus: Swiss Webster*<br>Wildtype | Charles River | Charles River Cat# NCI 551<br>IMSR Cat# TAC:sw,<br>RRID:IMSR_TAC:sw |
| Oligonucleotides |  |  |
| Recombinant DNA |  |  |
| Software and algorithms |  |  |
| limma | (Ritchie, et al., 2015) | LIMMA,<br>RRID:SCR_010943 |
| R statistical software | (Team, 2011) | R Project for Statistical Computing,<br>RRID:SCR_001905 |
| MACS version 2.1.0 | (Zhang, et al., 2008)<br>https://github.com/taoliu/MACS | MACS,<br>RRID:SCR_013291 |
| CisGenome | (Ji, et al., 2008) | CisGenome,<br>RRID:SCR_001558 |
| JASPAR motif database | (Khan, et al., 2018) | JASPAR,<br>RRID:SCR_003030 |
| Transfac motif database | (Matys, et al., 2006) | TRANSFAC,<br>RRID:SCR_005620 |
| Other |  |  |

#### Genomic Datasets

CUT&RUN, CUT&Tag, ATAC-Seq, ChIP-seq and RNA-seq datasets were deposited in GEO (GSE178838). The H3K27ac, H3K4me2 (GSE108880) and H3K4me1 ChIP-seq (GSE86690) datasets used in this study were processed and analyzed as all other ChIP experiments were done, described above. All chromosomal coordinates refer to the mm10 version of the mouse genome.

#### Embryonic manipulations

Experiments involving mice were approved by the Institutional Animal Care and Use Committee at the University of Texas at Austin (protocol AUP-2019-00233). The *Gli3^Xt-J^* and *Shh^tm1amc^* null alleles (Hui and Joyner 1993; Dassule et al. 2000) were maintained on a Swiss Webster background. The *Gli3^3XFLAG^* allele, with an N-terminal 3XFLAG-epitope, (Lopez-Rios et al. 2014; Lorberbaum et al. 2016) was maintained on a mixed background. For all genomic experiments, with the exception of *Shh^-/-^;Gli3^-/-^* ATAC samples which had to be individually genotypes, fresh E9.25 (21-23S) and E10.5 (32-35S) forelimb buds were pooled from multiple litters of the specified genotype. For ATAC-seq experiments with WT forelimbs (28-30S; 35S), and *Shh^-/-^* forelimbs (28-30S; 35S), posterior halves of forelimbs from 2-3 embryos were dissected and pooled. For RNA-seq, E9.25 whole forelimbs or E10.5 anterior halves forelimbs of WT and *Gli3^-/-^* forelimbs were pooled from 3-4 embryos; 3-4 biological replicates were used for each stage and genotype.

#### In situ hybridization

In situ hybridization done as previously described (Lewandowski et al., 2015). Fluorescent in situ hybridization was performed using HCR v3.0 Molecular Instruments reagents and was performed according to manufacturer’s protocol (Choi et al., 2018) except that probes were incubated overnight with a concentration of 16nM. Embryos were embedded in Ultralow gelling temperature agarose (Sigma A5030), cleared in Ce3D++ (Anderson et al., 2020) and visualized on a Zeiss LSM 710 confocal microscope and are shown as maximal intensity z-stack projections.

#### Immunofluorescence

Embryos were sucrose protected (30% sucrose) at RT and embedded in OCT. 12 μm cryosections were immediately fixed in 100% methanol for 3 minutes at −18C. Subsequently, sections were permeabilized in 0.1% PBST at RT for 15 min., blocked in 3% BSA (Fisher Scientific; BP1600-100), 5% Normal Goat Serum (Jackson Immunoresearch) in 0.1% PBST for 1 hr. at RT. Primary antibodies were incubated overnight with: 1:500 M2 FLAG (Sigma-Aldrich; F3165) and 1:500 Arl13b (Proteintech; 17711-1-AP). Secondary antibodies were incubated in block at RT for 1 hr. at the following dilutions: 1:500 Alexa568 (Thermo Fisher Scientific; A-11004) and 1:500 Alexa488 (Thermo Fisher Scientific; A-11034) and mounted with Prolong Gold mounting media (Thermo Scientific; P36930) prior to imaging.

#### Western Blots

Western blots were incubated for 1 hour at room temperature in 3% milk: 1:4000 M2 Flag (Sigma #F3165) and 1:2000 B-actin (Cell Signaling #8457). Secondary antibodies were incubated for 1 hours at room temperature in 3% milk: 1:5000 Donkey anti-mouse (Jackson #715-035-150), Donkey anti-rabbit (Jackson #711-005-0152). Blots were developed using Amersham ECL Prime (GE Healthcare #GERPN2232).

#### Chromatin Immunoprecipitation

ChIP-Seq was performed on whole E9.25 (21-23S) forelimbs pooled from 30-40 embryos as previously described (Vokes et al. 2008) with the following modifications. Cells were dissociated with 100μg/ml Liberase (Roche 05401119001) and fixed for 15 minutes in 1% formaldehyde. After cell lysis, chromatin was sheared in buffer containing 0.25% SDS with a Covaris S2 focused ultrasonicator. Antibodies used for ChIP experiments include H3K27ac (Abcam #ab4729) and H3K4me2 (Millipore #07-030). Libraries were generated using the NEBNext Ultra II library preparation kit with 15 cycles of PCR amplification (NEB E7645) and sequenced to a depth of >40 million reads per sample, using two biological replicates. Peaks were called using CisGenome version 2.1.0 (Ji et al. 2008). The read numbers were adjusted by library size and log2 transformed after adding a pseudo-count of 1. The differential analyses were performed using limma (Ritchie et al. 2015); peaks with an FDR < 0.05 were considered to have differential enrichment.

#### ATAC-Seq

WT E9.25 (21-23S) forelimbs or posterior forelimbs from E10 (28-30S) or E10.5 (35S) forelimbs were dissected and pooled from WT embryos (n=3, n=3). Posterior forelimbs from E10 (28-30S) or E10.5 (35S) were pooled from *Shh^-/-^* embryos (n=3, n=3). ATAC used components from the Nextera DNA Library Preparation Kit (Illumina) as described previously (Buenrostro et al. 2015) with the following variations. Limb buds were dissociated in 100μg/ml Liberase (Roche 05401119001) and lysed for 10 min. at 4°C, rotating, in a nuclear permeabilization buffer (5% BSA, 0.2% NP-40, 1mM DTT in Ca+/Mg+ free PBS + protease inhibitor) and then centrifuged for 10min. at 300g, 4C. Nuclei were resuspended, counted using Trypan Blue, and 30,000 nuclei from each WT and *Shh^-/-^* sample were put into each transposase reaction. The transposase reaction (30uL volume) was carried out for 1 hour at 37°C with gentle agitation. Libraries were generated using 11 cycles of PCR amplification with NEB high fidelity 2x master mix (New England Biolabs), cleaned up with SparQ PureMag beads (QuantaBio) and sequenced on an Illumina NovaSeq using PE150 or SR100 to a depth of at least 40 million reads. Peaks were called using MACS2 with a fixed window size of 200bp and a q-value cutoff of 0.05. Read counts from the peak regions were obtained and differential analysis was performed using DESeq2 (Love et al., 2014). Peaks with FDR < 0.05 in the differential test were considered to have significant chromatin accessibility difference.

Footprinting analysis was performed using HINT (Li et al., 2019) based on the ATAC-seq data. After obtaining the DNA footprints, GLI3 and HAND2 motifs were mapped to the footprints which overlap with regions that have pre-HH only GLI3 binding, E10.5 HAND2 binding, and E9.25 H3K27ac signals (Figure S6C) using CisGenome (Ji et al., 2008). Motif enrichment (Figure S6E) is calculated as the ratio between the percentage of footprints containing the motif in the E9.25 WT samples and that in the E10.5 *Shh^-/-^* samples. Fisher’s extract test is applied to test if the motif enrichment is different between E9.25 WT and E10.5 *Shh^-/-^* (GLI3 p-value = 0.87, HAND2 p-value = 0.14).

#### CUT&Tag

Experiments were performed as described in (Kaya-Okur et al., 2019) with the following modifications. 100,000 cells from E9.25 (21-23S) forelimbs were dissociated, bound to Concanavalin beads and incubated overnight at 4°C on a nutator with primary antibodies. Antibodies were used at the following concentrations: 1mg/mL of H3K4me1 (Abcam #ab8895) and 5mg/mL of H3K27me3 (abcam #ab195477). The following day, samples were incubated at room temperature for 30 min. with secondary antibody, 1:500 Donkey anti-rabbit (Jackson ImmunoResearch #711-005-152). CUT&Tag transposases used were gifted from the Henikoff lab and EpiCyphr (now commercially available). Libraries were generated using NEB high fidelity 2x master mix with 14 PCR cycles and cleaned up to remove adapters using or SparQ PureMag beads (QuantaBio). Samples were sequenced on an Illumina NextSeq 500 instrument using PEx75 to a depth of depth of 3-5 million reads. Peaks were called using SEACR (Meers et al., 2019).

#### CUT&RUN

Experiments were performed according to EpiCypher’s CUTANA CUT&RUN protocol (EpiCypher #15-1016) with the following modifications. 200,000-300,000 cells from E9.25 (21-23S) forelimbs and anterior E10.5 (32-35S) forelimbs were incubated overnight at 4°C on a nutator with 1:250 FLAG primary antibody (Sigma #F3165), 1:100 HDAC1 (Abcam ab7028) or 1:200 HDAC2 (Abcam ab7029). The following day, samples were incubated at room temperature for 30 min. with secondary antibody, 1:500 Donkey anti-mouse (Jackson ImmunoResearch #715-035-150) or Donkey anti-rabbit (Jackson ImmunoResearch #711-005-152), followed by three washes in Digitonin wash buffer. CUTANA pAG-MNase was then incubated with samples for 10 minutes at room temperature and then the MNase reaction was performed for 2 hours at 4°C on a nutator. Libraries were generated using NEBNext Ultra II DNA Library Prep Kit with 14 PCR cycles and cleaned up to remove adapters using AMPure XP beads (Beckman Coulter) or SparQ PureMag beads (QuantaBio). Samples were sequenced on an Illumina NextSeq 500 instrument using PEx75 to a depth of depth of 3-5 million reads. Peaks were called using MACS (Zhang et al., 2008).

#### Intron retention rates

The formula (1-N)/M was used to calculate the intron retention rate for each gene, where N= the number of reads overlapping with any of the gene’s exons and M= the number of reads overlapping with the body of the gene. Limma (Ritchie et al., 2015) was used to identify differential intron retention rate between E9.25 and E10.5 *Gli3^-/-^* limbs, FDR cutoff< 0.05.

#### Identification of GLI target genes

To identify GLI target genes, we developed a weighted system which incorporated HH-responsive GBRs, poised chromatin modifications, active promoter marks and genes identified as being significantly downregulated in E10.5 *Shh^-/-^* RNA-seq compared to WT limb buds (FDR=0.05; WT/*Shh^-/-^* FC <0) (Lewandowski et al., 2015; GSE58222). HH-responsive GBRs were intersected with poised enhancer modifications from H3K4me1 ChIP-seq (GSE86690), H3K4me2 ChIP-seq and E10.5 ATAC-seq datasets (Lex et al., 2020; GSE108880) from E10.5 WT limb buds. HH-responsive GBRs enriched for poised enhancers modifications, and thus increasing the probably of being active enhancers, were given a higher weight than ones without modifications. Genes identified as downregulated in *Shh* mutant limbs were intersected with H3K4me3 (GSE86698) and H3K4me6 (GSE8669) E10.5 WT limb datasets. Genes enriched for these active promoter marks were given more weight. The weighted HH-responsive GBRs and weighted genes were then intersected based on proximity, up to 500kb, and within the same topologically associated domain (TAD) (GSE96107). Genes were identified as a putative HH targets based on their proximity to HH-responsive GBRs and the number of HH-responsive GBRs, such that a gene with multiple HH-responsive GBRs, enriched for poised enhancer marks, would rank higher than a gene with HH-responsive GBRs found more distally. https://github.com/Boksunni/Predicted-HH-List-Generation

## Supplemental Tables

**Supplemental Table 1.** Predicted direct GLI3 target genes (tab 1). Predicted direct target genes enriched with H3K27me3 and H3K27ac in pre-HH limb buds (tab 2).

## Supplemental Data Files

**Supplemental File 1.** Called peaks for CUT&RUN and CUT&Tag datasets. Includes: E9.25 and anterior E10.5 GLI3 CUT&RUN, E9.25 H3K4me1 CUT&Tag, E9.25 H3K27me3 CUT&Tag, E10.5 anterior and posterior H3K27me3 CUT&Tag, E9.25 HDAC1, E9.25 HDAC2, E10.5 HDAC1, E10.5 HDAC2.

**Supplemental File 2.** Differential Chip-seq analyses. Differential analysis of H3K4me2 ChIP-seq enrichment in E9.25 (21-23S) and E10.5 (32-35S) WT limb buds (tab 1). Differential analysis of H3K27ac ChIP-seq enrichment in E9.25 (21-23S) vs. E10.5 (32-35S) WT limb buds (tab 2) and E9.25 (21-23S) WT vs. *Gli3^-/-^* limb buds (tab 3).

**Supplemental File 3.** Differentially expressed genes in RNA-seq datasets. Differentially expressed genes in WT vs. *Gli3^-/-^* E9.25 limb buds (tab 1), WT vs. *Gli3^-/-^* E10.5 anterior limb buds (tab 2) and E9.25 vs. E10.5 anterior WT limb buds (tab3). Intron retention rates in E9.25 *Gli3^-/-^* limb buds and E10.5 *Gli3^-/-^* limb buds (tab 3). ‘IRtype’ denotes if a gene has a significantly higher IR rate at E9.25, E10.5 or neither (labeled ‘other’).

**Supplemental File 4.** Differential ATAC-seq comparing E9.25 WT whole limb buds, E10 WT (28-30S), E10 *Shh^-/-^* (28-30S), E10.5 WT (35S) and E10.5 *Shh^-/-^* posterior limb buds.

---

**Figure S1.**
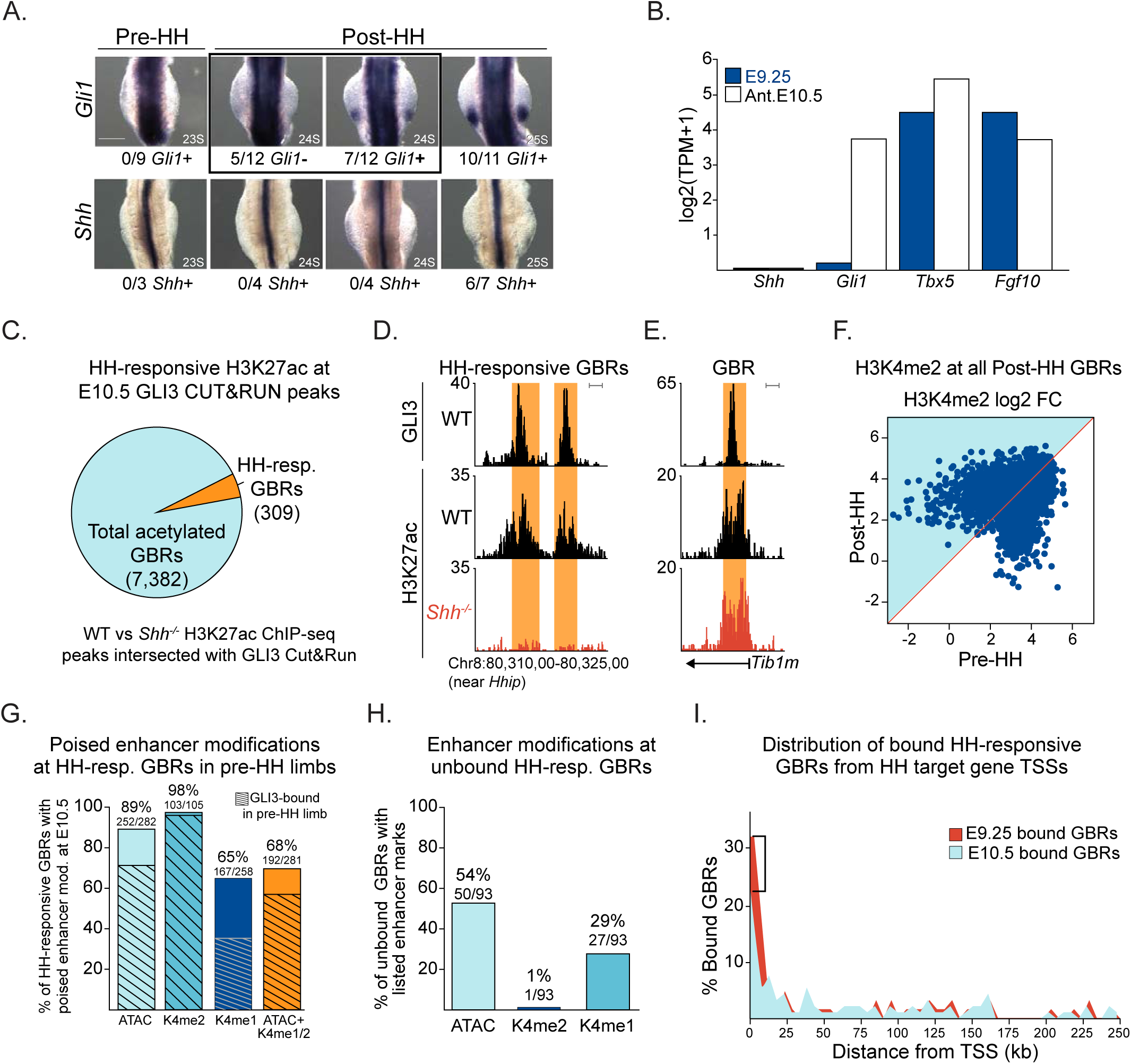
Properties of GLI3 binding at enhancers before and after the onset of HH signaling. A. Whole mount in situ hybridization for *Gli1* and *Shh* in 23-25S embryos to define the onset of HH induction. While *Shh* is not detectable in the limb until 25S, the earliest detection of *Gli1* is at 24S, where 5/15 embryos have *Gli1* expression. B. Expression values for *Shh, Gli1* and selected limb genes, in E9.25 and E10.5 anterior limb buds from RNA-seq data. C. Intersection of E10.5 GLI3 CUT&RUN peaks with previously published WT vs *Shh^-/-^* E10.5 H3K27ac ChIP-seq peaks (Lex et al., 2020). H3K27ac peaks reduced in *Shh^-/-^* limbs that overlap GLI3 binding regions were categorized as HH-responsive GBRs. D, E. Examples of a HH-responsive GBR that loses acetylation in *Shh^-/-^* limbs (D) and a GBR that maintains acetylation in the absence of HH signaling (E). Orange shading in tracks indicate the GBRs. F. Scatterplot showing H3K4me2 ChIP-seq fold enrichment at E10.5 GBRs in pre-HH (21-23S) and post-HH (32-35S) limb buds (n=2 biological replicates at each stage). No peaks were significantly changed between pre- and post-HH signaling. G. Percentage of HH-responsive GBRs enriched (called peaks) for the poised enhancer markers H3K4me1 (CUT&Tag, n=3) and H3K4me2 (n=2) and accessible chromatin (ATAC-seq peaks, n=2) prior to HH signaling. Percentages indicate the number of E9.25 GBRs enriched for the specified feature, out of the total number of HH-responsive GBRs enriched for that enhancer modification at E10.5. H. Enhancer modifications present at HH-responsive GBRs that are not bound by GLI3 at E9.25. I. Distribution of E9.25 and E10.5 GBRs from the TSSs of putative HH target genes. Scale bars for tracks indicate 1kb.

**Figure S2.**
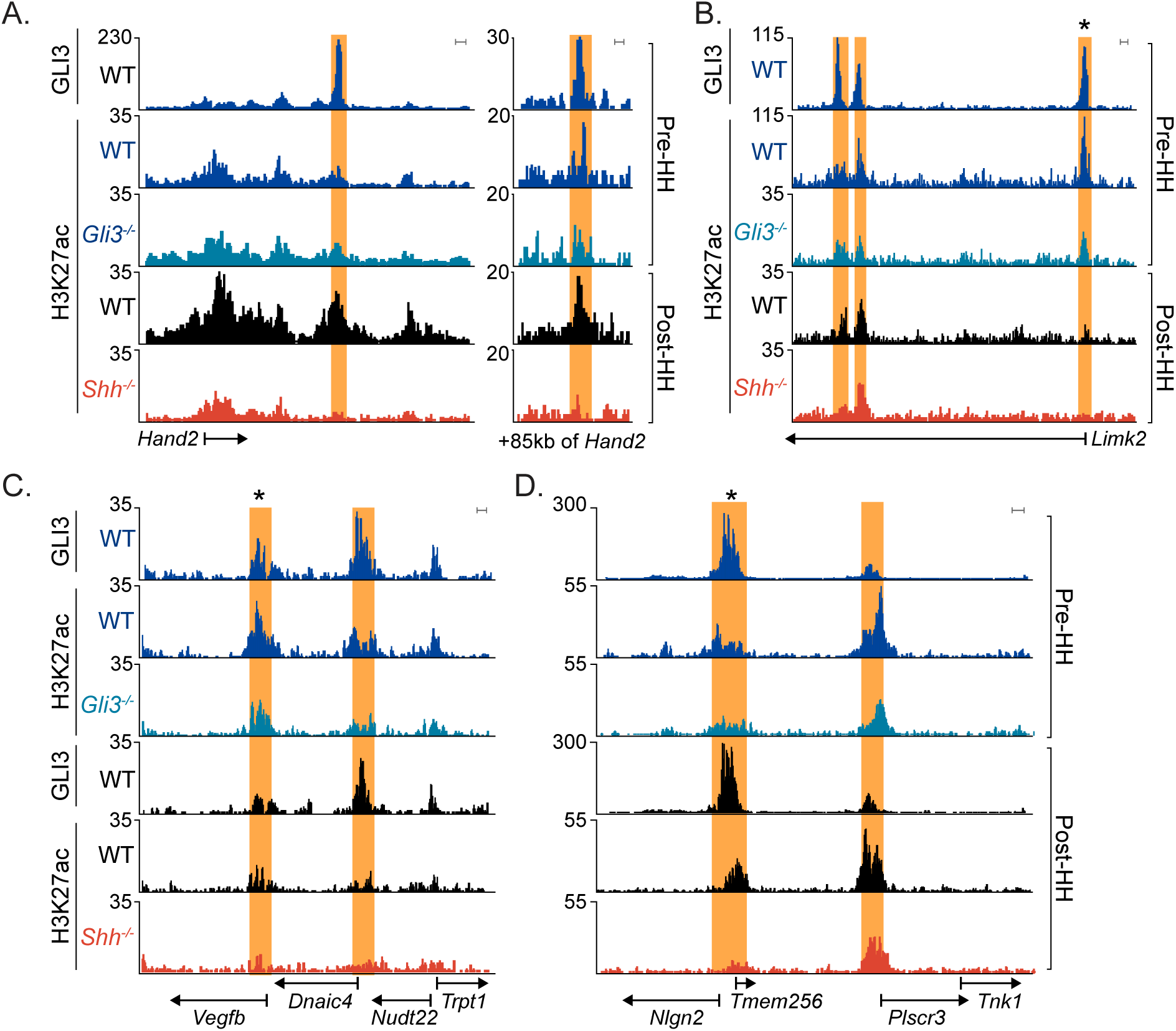
E9.25 acetylated GBRs that do not further increase H3K27ac levels with loss of *Gli3*. A. HH-responsive GBRs (orange shading) near the GLI3 target Hand2 that have H3K27ac enrichment at E9.25 do not have increased levels of H3K27ac in *Gli3^-/-^* limb buds, suggesting there is no de-repression of *Hand2*. B-D. HH-responsive GBRs (orange shading) with significantly higher acetylation prior to HH induction (indicated by asterisk; FDR<0.05), do not have increased acetylation with the absence of *Gli3*. Scale bars for tracks indicate 1kb.

**Figure S3.**
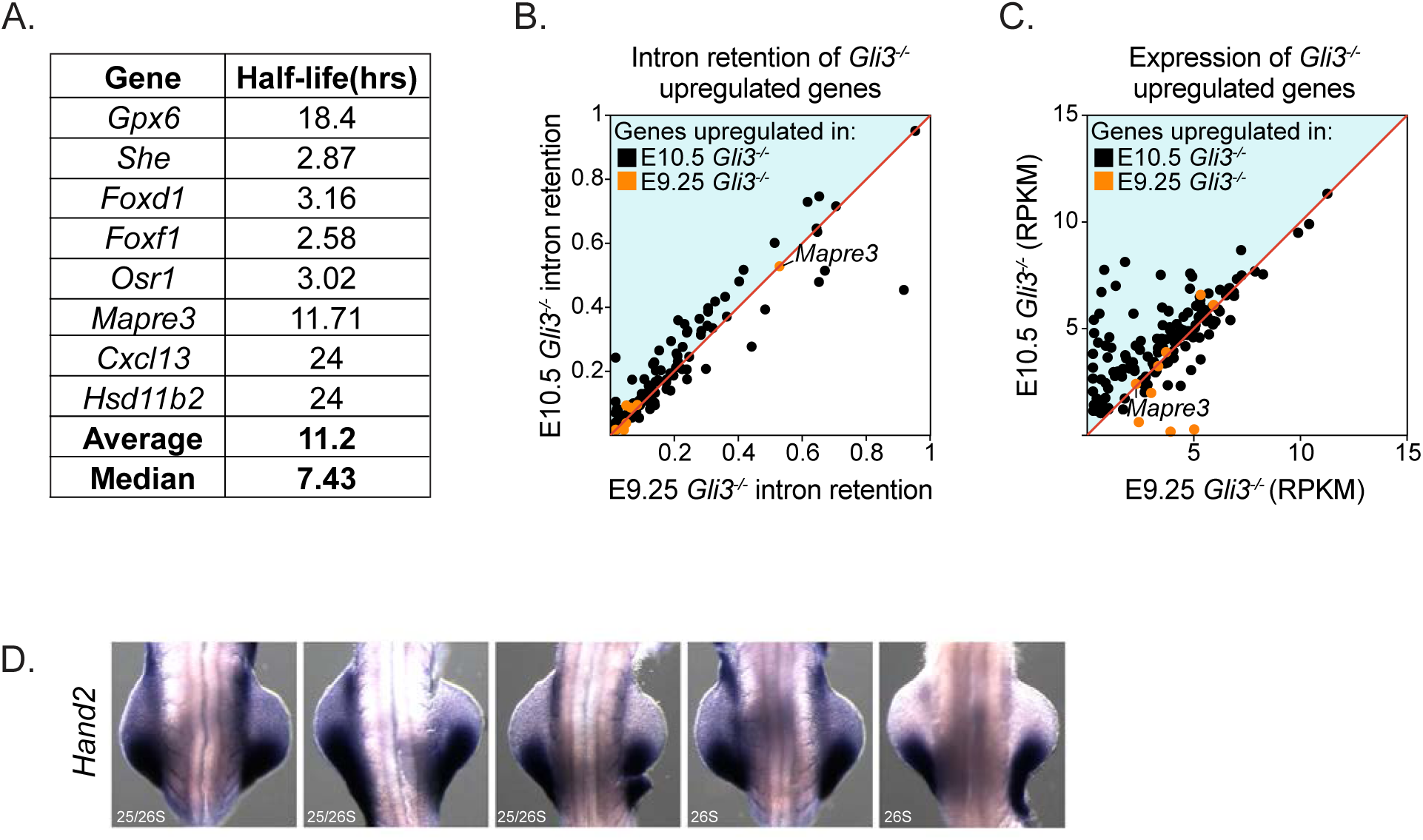
E9.25 *Gli3^-/-^* upregulated genes are expressed at low levels and have low intron retention rates. A. Table of half-lives E9.25 Gli3^-/-^ upregulated genes, previously determined in differentiating embryonic stem cells (Sharova et al., 2009). B. Intron retention levels in E9.25 and E10.5 *Gli3^-/-^* upregulated genes. While E10.5 upregulated genes have intron retention rates that vary, E9.25 upregulated genes have intron retention rates near 0, suggestive of mature transcripts. The exception to this is one gene, *Mapre3*, which has high intron retention in both E9.25 and E10.5 *Gli3^-/-^*, but is not upregulated at E10.5. The formula (1-N)/M was used to calculate the intron retention rate for each gene, where N= the number of reads overlapping with any of the gene’s exons and M= the number of reads overlapping with the body of the gene. Limma (Ritchie et al., 2015) was used to identify differential intron retention rate between E9.25 and E10.5 *Gli3^-/-^* limbs, FDR cutoff< 0.05. C. Expression of genes upregulated in E9.25 and E10.5 *Gli3^-/-^* limb buds. D. Expression of Hand2 in WT limb buds at ∼26S. Note that most embryos have some anterior expression of Hand2 and Hand2 is only completely posteriorly restricted in one embryo at this stage (far right).

**Figure S4.**
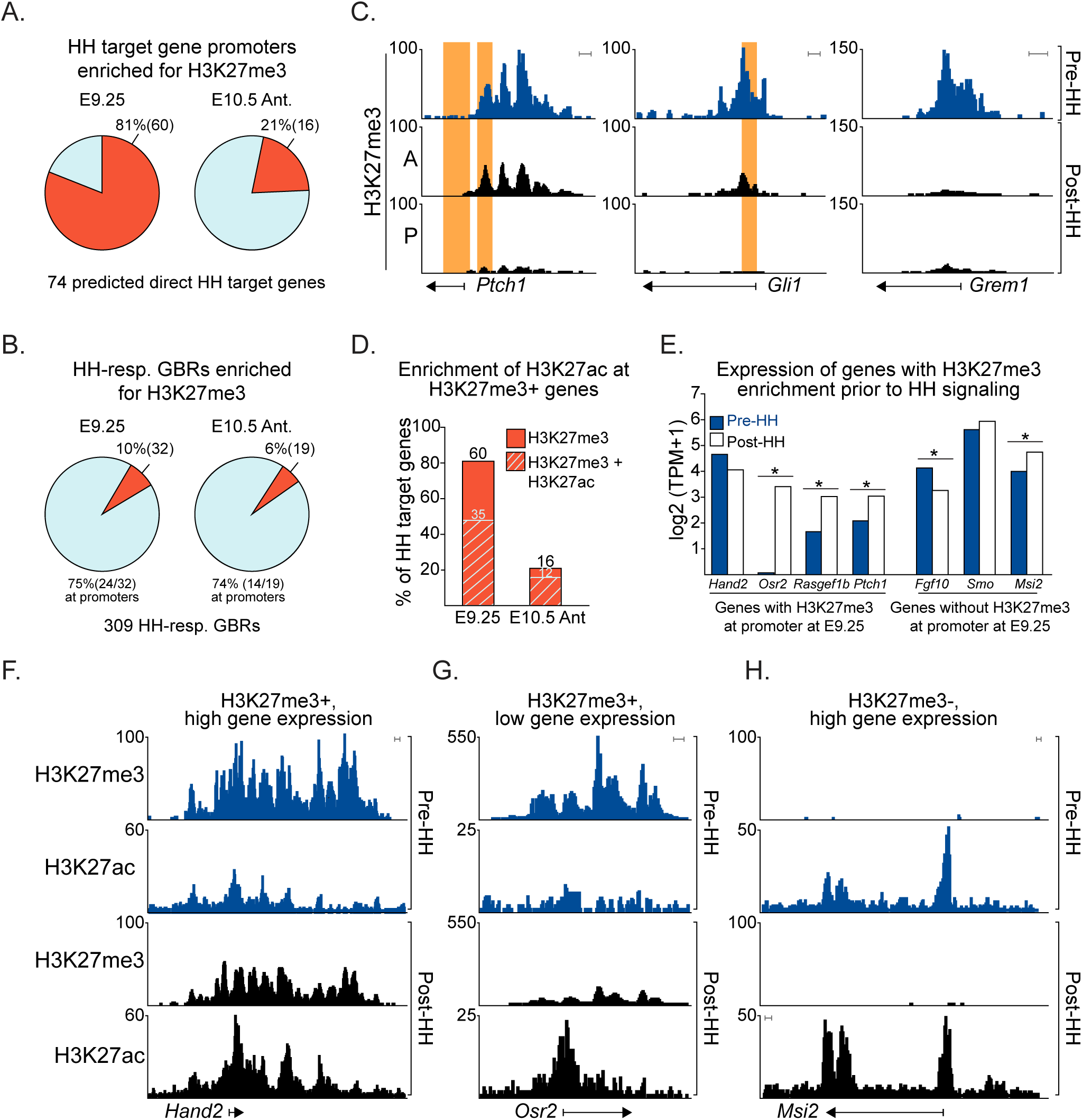
H3K27me3 is enriched at many HH target genes prior to HH signaling. A,B. H3K27me3 enrichment at the promoters of predicted direct HH target gene promoters (A) and HH-responsive GBRs (B) in E9.25 and E10.5 anterior limb buds. C. Examples of H3K27me3 enrichment at promoters of HH target genes in E9.25 and anterior and posterior E10.5 limb buds. Orange shading indicates the binding regions for HH-responsive GBRs defined in Figure S1C. D. Percentage of HH target gene promoters with H3K27me3 and H3K27ac at E9.25 and E10.5. E. Relative expression levels of genes at E9.25 and E10.5 derived from RNA-seq data (Supplemental File 3) with and without H3K27me3 enrichment at promoters at E9.25. F-H. Examples of H3K27me3 and H3K27ac enrichment of genes with high expression at E9.25 with H3K27me3 enrichment (F), low expression with H3K27me3 enrichment (G) and high expression without H3K27me3 enrichment (H). Scale bars for tracks indicate 1kb.

**Figure S5.**
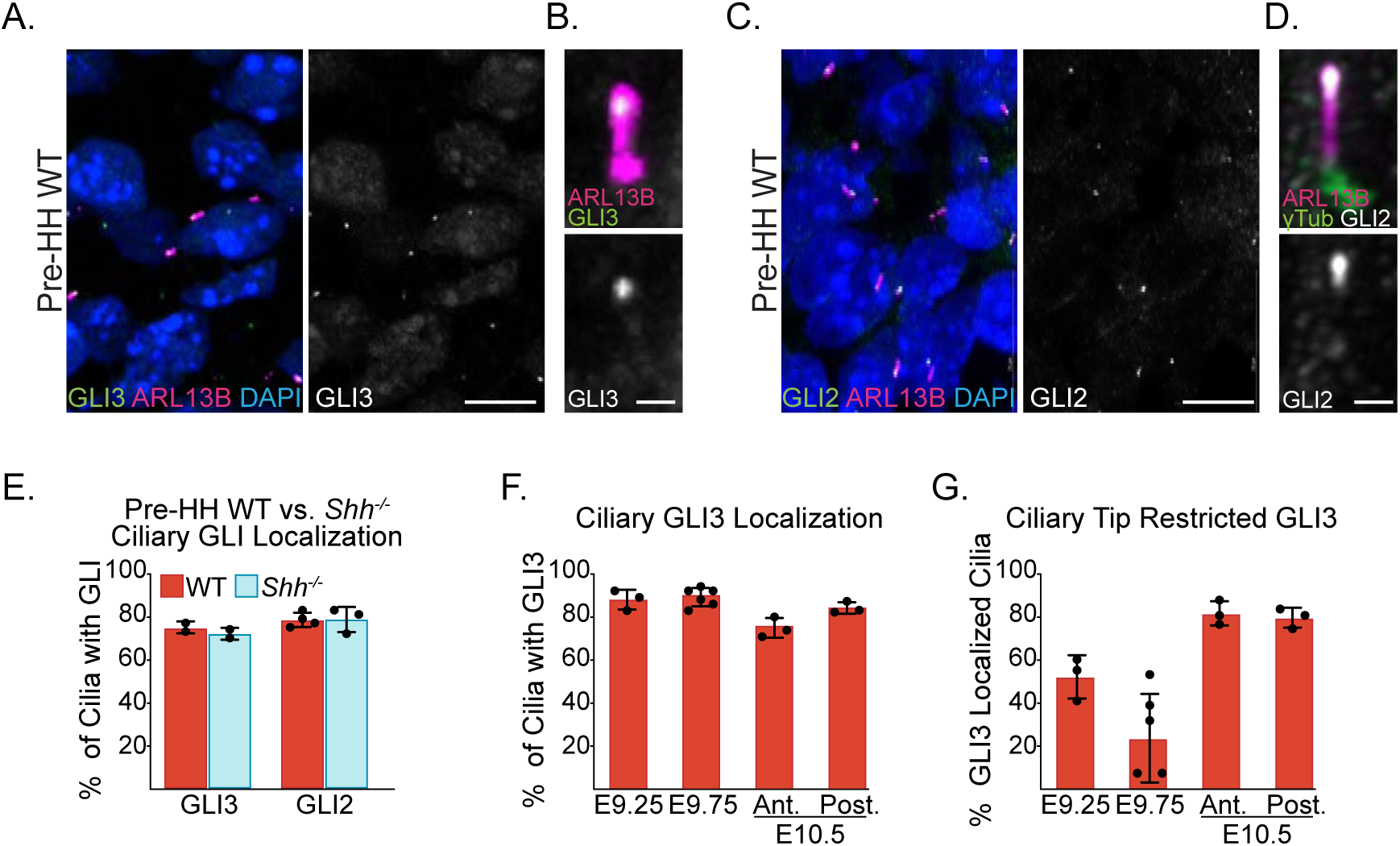
GLI ciliary distribution and localization in developing limbs. Images were collected with a Nikon A1R Resonant Scanning Confocal System and are maximum intensity projections. A,B. Endogenous GLI3^FLAG^ and ARL13b in a representative pre-HH E9.25 WT limb bud, and a representative cilium with GLI3^FLAG^ localization. C,D. Endogenous GLI2, ARL3b and {g-TUBULIN (basal body) in a representative pre-HH E9.25 WT limb bud. GLI2 is localized to the opposite ciliary end as γ-TUBULIN indicating most GLI localization is at the ciliary tips and not the base (D). E. Quantification of GLI2 and GLI3^FLAG^ ciliary localization in E9.25 WT and *Shh^-/-^* limb buds (n=2 and 4 biological replicates, respectively). Note the similar levels of GLI2 and GLI3 ciliary localization in both genotypes, suggesting unprocessed GLI proteins are localized at the ciliary tip. F-G. Quantifications of GLI3 ciliary localization (D) in pre-HH E9.25 (21-23S; n=3), E9.75 (26-28S n=6) distal limb buds and E10.5 (35S; n=3) anterior and posterior limb buds. E. Quantification of cilia in the same dataset with GLI3 restricted to the ciliary tip. Error bars in E-G indicate SEM. Scale bars for panels A and C indicate 10µm; scale bards for panels B and D indicate 0.5µm.

**Figure S6.**
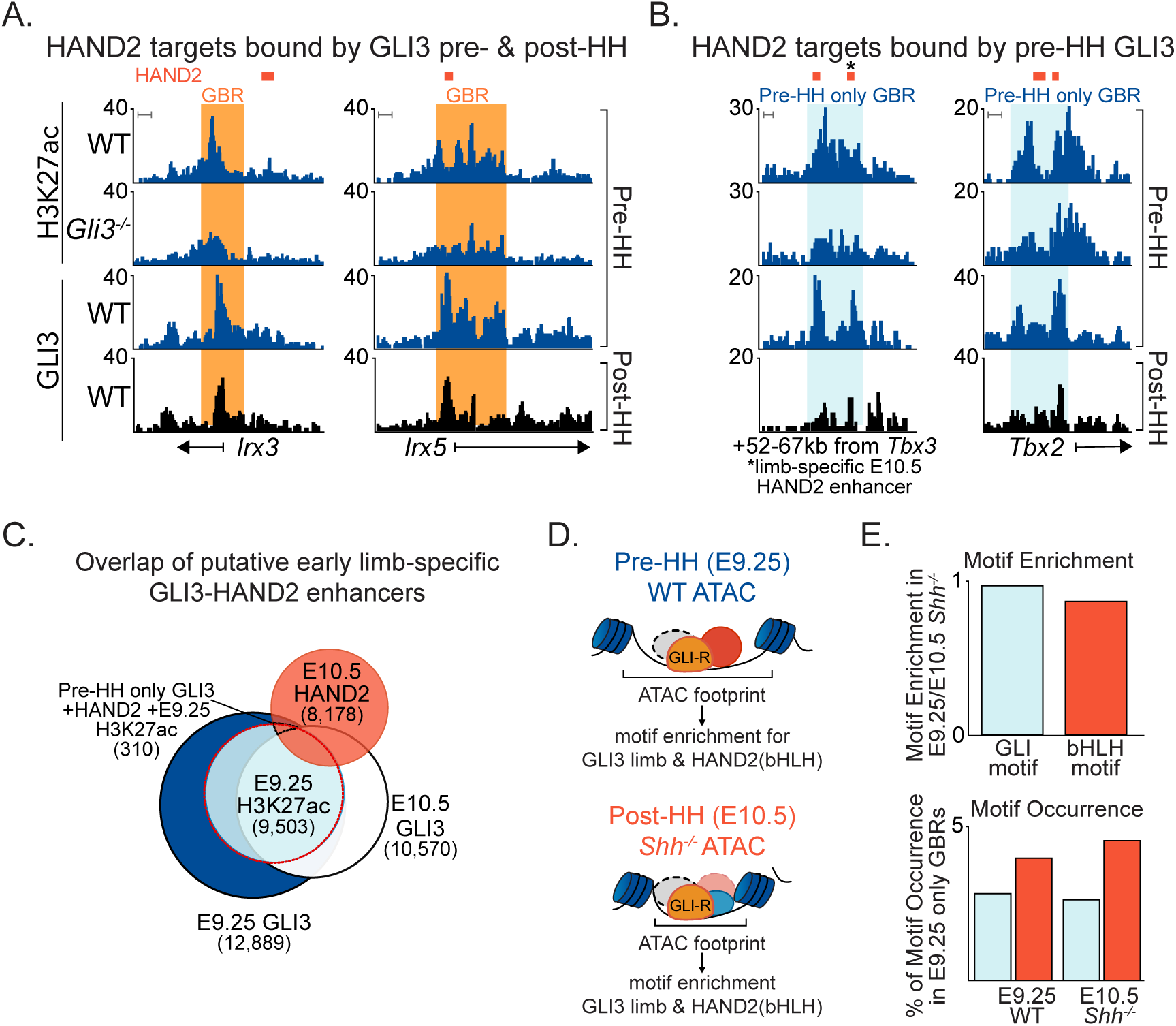
Co-localization of HAND2 and GLI3 in pre-HH limb buds. A. Examples of genes with HAND2 binding regions in E10.5 limbs (red bars denote *Hand2*regions identified in Osterwalder et al, 2014) that are also bound by GLI3 (orange shading) in pre-HH or post-HH E10.5 limb buds. B. HAND2 target genes *Tbx2* and *Tbx3* have HAND2 bound regions that overlap with pre-HH specific GBRs (blue shading) but not post-HH GLI3 binding regions. Asterisks denote previously identified HAND2-bound enhancers (Osterwalder et al., 2014). C. Venn diagram of pre- and post-HH (E10.5) GLI3 binding regions, pre-HH H3K27ac peaks and previously identified E10.5 HAND2 limb binding sites (Osterwalder et al., 2014). 310 regions are bound by GLI3 only in pre-HH limb buds (not post-HH), overlap with HAND2 binding regions and are acetylated, indicating possibility of these regions being active enhancers. D. Schematic for performing motif enrichment in ATAC footprints identified in pre-HH WT limbs and compared to ATAC footprints in E10.5 *Shh^-/-^* limbs. E. Quantification of enrichment (top) and occurrence (bottom) of limb GLI3 and face HAND2 motifs in the 310 acetylated pre-HH only GBRs overlapping HAND2 binding sites, compared to all acetylated pre-HH GBRs (red circle in D), for both pre-HH WT and post-HH E10.5 *Shh^-/-^* limbs (Lex et al., 2020; Osterwalder et al., 2014). Note that GLI3 and HAND2 motif enrichment and occurrence are unchanged pre-HH WT and post-HH E10.5 *Shh^-/-^* limbs. Scale bars denote 1kb.

